# Extraocular muscle stem cells exhibit distinct cellular properties associated with non-muscle molecular signatures

**DOI:** 10.1101/2023.03.10.532049

**Authors:** Daniela Di Girolamo, Maria Benavente-Diaz, Alexandre Grimaldi, Priscilla Thomas Lopes, Melania Murolo, Brendan Evano, Stamatia Gioftsidi, Vincent Laville, Sebastian Mella, Shahragim Tajbakhsh, Glenda Comai

## Abstract

The muscle stem cell (MuSC) population is recognized as functionally heterogeneous. Cranial muscle stem cells, which originate from head mesoderm, can have greater proliferative capacity in culture and higher regenerative potential in transplantation assays when compared to those in the limb. The existence of such functional differences in phenotypic outputs remain unresolved as a comprehensive understanding of the underlying mechanisms is lacking. We addressed this issue using a combination of clonal analysis, live imaging, and scRNA-seq, identifying critical biological features that distinguish extraocular (EOM) and limb (Tibialis anterior, TA) MuSC populations. Time-lapse studies using a *Myogenin^tdTomato^*reporter showed that the increased proliferation capacity of EOM MuSCs is accompanied by a differentiation delay *in vitro*. Unexpectedly, in vitro activated EOM MuSCs expressed a large array of distinct extracellular matrix (ECM) components, growth factors, and signaling molecules that are typically associated with mesenchymal non-muscle cells. These unique features are regulated by a specific set of transcription factors that constitute a coregulating module. This transcription factor network, which includes Foxc1 as one of the major players, appears to be hardwired to EOM identity as it is present in quiescent adult MuSCs, in the activated counterparts during growth and retained upon passages in vitro. These findings provide insights into how high-performing MuSCs regulate myogenic commitment by active remodeling of their local environment.

## INTRODUCTION

Skeletal muscle stem cells are arguably the most diverse in terms of molecular properties compared to other adult stem cells that are distributed throughout the body. Genetic and transcriptomic studies have shown that the MuSC population in any particular anatomical location is heterogeneous. Certain subsets are more prone to self-renewal or differentiation, differ in transplantation efficiency, as well as their stem cell-niche interactions, metabolism, and resistance to stress upon activation (Barruet et al., 2020; Chakkalakal et al., 2014; Cho et al., 2020; Dell’Orso et al., 2019; Dumont et al., 2015; Gayraud-Morel et al., 2012; Hernando-Herraez et al., 2019; Micheli et al., 2020; Morree et al., 2019; Ono et al., 2012; Rocheteau et al., 2012; Scaramozza et al., 2019; Tierney et al., 2018; Vartanian et al., 2019; Yartseva et al., 2020; Yennek et al., 2014). Despite this diversity, MuSCs share common functions as they are essential for postnatal growth and repair of the skeletal muscle system (Lepper, 2011; Murphy, 2011; Sambasivan, 2011). In response to muscle tissue damage or growth factors in culture media, MuSCs activate, proliferate and undergo cell fate decisions to either self-renew to replenish the MuSC pool (expressing *Pax7)*, or commit to the myogenic program and differentiate into myoblasts and fusion-competent myocytes that express the commitment and differentiation factors *Myod* and *Myog*, respectively (Evano and Tajbakhsh, 2018; Zammit et al., 2006).

An unexpected finding was the discovery that MuSCs in different anatomical locations are programmed with distinct upstream transcription factors (TF) prior to acquiring myogenic identity (Gopalakrishnan et al., 2015; Harel et al., 2009; Kelly et al., 2004; Sambasivan et al., 2009; Tajbakhsh, 1996). This intriguing observation correlates with reports where subsets of cranial MuSCs are functionally more robust in terms of proliferation, engraftment efficiency, and susceptibility to disease, when compared to those in the limb (Randolph and Pavlath, 2015). However, the intrinsic and extrinsic regulators conferring robust features to cranial muscles have not been characterized.

EOMs are a subset of head muscles that govern eye movements. EOMs are derived from unsegmented cranial mesoderm and are regulated by distinct transcriptional factors and signalling molecules compared to the somite-derived limb and trunk muscle groups (Grimaldi and Tajbakhsh, 2021; Michailovici et al., 2015; Sambasivan et al., 2011). Notably, when the transcription factor *Pax3* is inactivated, no limb muscles develop, yet cranial muscles including the EOMs are present (Sambasivan et al., 2009; Tajbakhsh et al., 1997). Conversely, mice lacking the transcription factor *Pitx2* do not form EOMs whereas other cranial and somite derived muscles are unaffected (Diehl et al., 2006; Gage et al., 1999; Zacharias et al., 2011). Additionally, whole muscle transcriptional profiling revealed an extraordinary diversity amongst adult skeletal muscle groups (Kui et al., 2022; Terry et al., 2018). As some muscle subsets, like the EOMs, are preferentially spared in muscular dystrophies and during ageing (Emery, 2002; Formicola et al., 2014; Verma et al., 2017), it has been proposed that intrinsic properties of MuSCs or the respective myofibers could determine their differentially sensitivity to disease (Randolph and Pavlath, 2015; Terry et al., 2018).

Similar to MuSCs derived from other craniofacial muscles (Ono et al., 2010; Terry et al., 2018), EOM MuSCs have a greater proliferative capacity in culture when compared to those in the limb, with this property being conserved in *Mdx* mice, a model of Duchenne muscular dystrophy (Dmd), and in a preclinical rat model of the disease (Stuelsatz et al., 2015; Taglietti et al., 2023). Additionally, EOM MuSCs have a higher engraftment potential compared to those in the limb (Stuelsatz et al., 2015). Although debated (McLoon and Wirtschafter, 2002; McLoon and Wirtschafter, 2003; Stuelsatz et al., 2015; Verma et al., 2017), EOM MuSCs were reported to continuously contribute to myofibres in homeostatic conditions, suggesting a higher level of basal activation of the MuSC population (Keefe et al., 2015; Pawlikowski et al., 2015; Taglietti et al., 2023). In addition, changes in the transcriptomic profile of EOM MuSCs after transplantation into limb muscle (Evano et al., 2020) showed that despite a large transcriptional reprogramming after transplantation, about 10% of EOM specific genes persisted in the heterotypic niche environment. This finding suggests that cell autonomous regulation of MuSC properties predominates to a certain extent in EOM MuSCs (Evano et al., 2020). Moreover, recent single-cell transcriptomic analysis identified the thyroid hormone signalling pathway as a key factor preventing entry into senescence of EOM MuSCs in Dmd rats (Taglietti et al., 2023). Yet, whether deeply-rooted TF gene regulatory networks exist within MuSCs subsets, and contribute to the maintenance of anatomically-distinct MuSC phenotypes remains largely unexplored.

Here, we used droplet-based scRNA-seq to investigate the transcriptional states that govern the outperformance of MuSC subsets during activation. By integrating genome-wide analysis with cell biology approaches using mouse reporter lines, we identified key regulators that confer distinct mesenchymal-like features to activated EOM MuSCs and expression of extracellular matrix (ECM) components. Computational analyses showed that this pool of EOM cells displays features of a more stem-like state that is actively maintained during *in vitro* proliferation through a specific set of transcription factors forming a co-regulatory module.

## RESULTS

### Functional differences among MuSCs are accompanied by heterogeneity in myogenic gene expression dynamics

In agreement with previous reports (Stuelsatz et al., 2015), we show that culture of EOM and *Tibialis anterior* (TA) MuSCs isolated by FACS using *Tg:Pax7-nGFP* mice (Sambasivan et al., 2009), revealed a larger fraction of proliferative cells in the EOM from day 3 (D3) onwards, peaking at D4 and remaining higher even when differentiation and fusion starts to take place at D5 (**Figure 1A-C**). Cell proliferation, monitored by 5-Ethynyl-2’-deoxyuridine (EdU) uptake, revealed 45,1% of Pax7+EdU+ EOM cells, compared to 19,8% in the TA at D5 (**Figure 1B**). To obtain insights into the cellular basis for the outperformance of MuSC subsets upon exit from quiescence, we performed clonal and live imaging analysis *in vitro*. First, MuSCs from EOM and TA were plated at clonal density in 96-well plates. In parallel, a fraction of the isolated cells was plated as a bulk culture, allowed to activate for 2 days *in vitro*, re-isolated and then plated at clonal density (**Suppl Figure 1A**). The total cellular output in both cases was quantified two weeks after seeding. In accordance with a previous study (Stuelsatz et al., 2015), EOM MuSCs displayed a markedly higher clonal capacity than those from the TA muscle, with a mean of 1315 cells/clone and 186 cells/clone respectively (7-fold difference; **Suppl Figure 1B,** T1). Surprisingly, the clonogenic properties of EOM MuSCs continued to be elevated after 2 days of activation *in vitro* where they yielded 3656 cells/clone compared to 297 cells/clone for activated TA MuSCs (12-fold difference; **Suppl Figure 1B**, T2).

**Figure 1.**
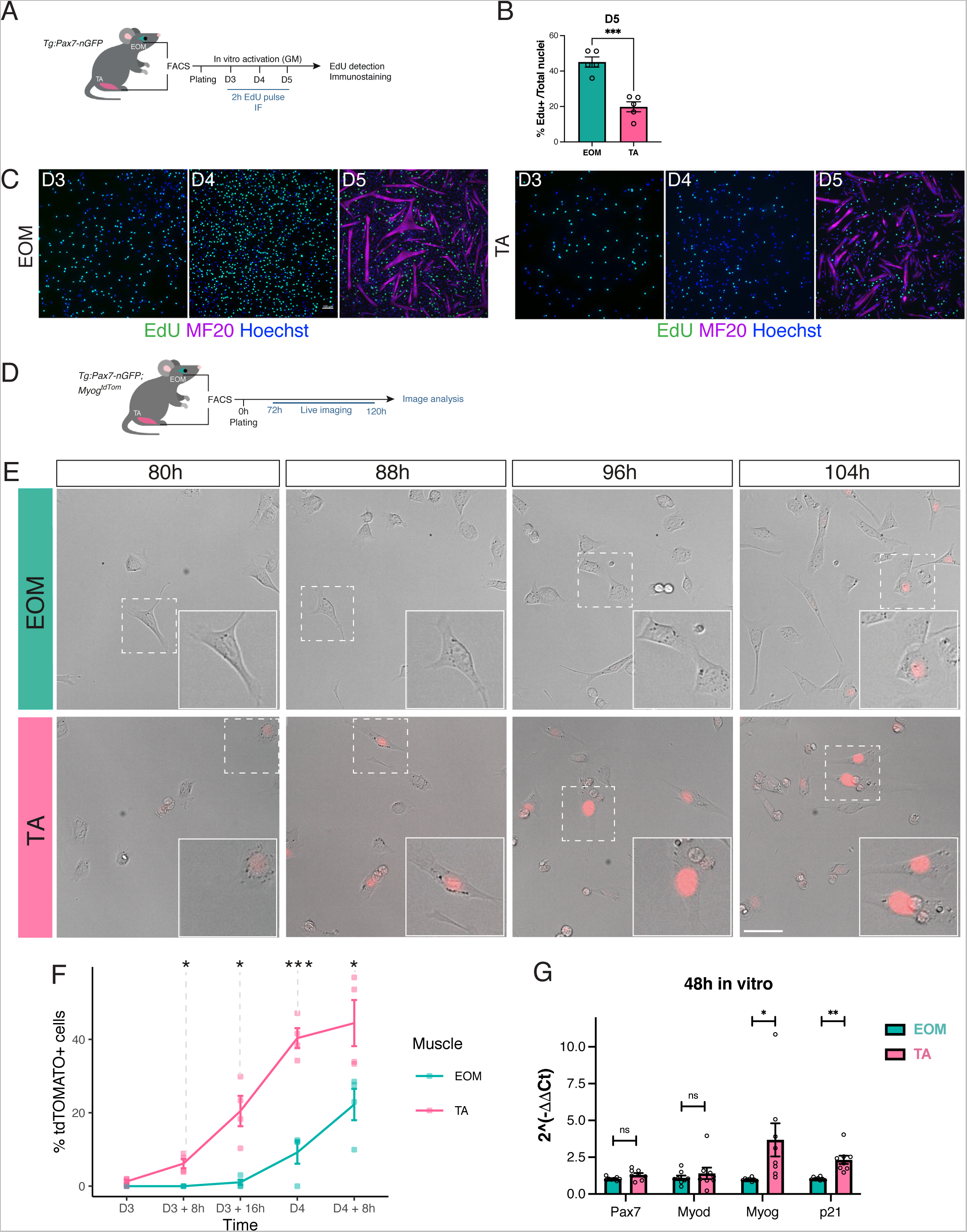
EOM and TA MuSCs exhibit functional differences following activation. A. Experimental scheme. MuSCs were isolated by FACS based on GFP fluorescence from *Tg:Pax7-nGFP* mice, plated and pulsed with EdU before fixation and immunostaining at Day (D) 3, D4 and D5. B. Quantification of the percentage of EdU+ cells at D5. Two-tailed unpaired Student’s t-test. *** p-value<0.005. n=5 mice. C. Immunofluorescence for MF20 and EdU detection at different time points of the experiment in A. D. Experimental scheme. EOM and TA MuSCs from *Tg:Pax7-nGFP;Myog^tdTom^* mice were cultured for 72h and imaged for 48h. E. Representative overlayed DIC and red fluorescence channel images at selected time points showing a temporal increase of tdTOMATO fluorescence. Scale bar 25 μm. F. Quantification of selected timepoints in (E). Graph represents mean percentage of tdTOMATO^+^ cells over total number of cells at each timepoint for each muscle. Percentage of tdTOMATO^+^ cells was compared between EOM and TA. Two-tailed unpaired Student’s t-test; * p-value<0.05, *** p-value<0.005. n=4 mice; > 100 cells counted. G. EOM and TA MuSCs were cultured for 48h and then processed for RNA extraction. RT-qPCR for *Pax7*, *Myod*, *Myogenin* and *p21* normalized to *Rpl13*. Two-tailed unpaired Student’s t-test; *p-value<0.05, *** p-value<0.01. n=8 mice.

We then monitored their differentiation dynamics *in vitro* by live imaging using the *Tg:Pax7-nGFP;Myog^ntdTom^* mouse lines (Benavente-Diaz et al., 2021; Sambasivan et al., 2009) to isolate MuSCs by FACS and then monitor the onset of *Myog* expression using a nuclear localised tdTOMATO fluorescent reporter. After activation for 3 days *in vitro*, EOM and TA cells were imaged every 12 min for ∼48h (**Figure 1D**). Individual cells were tracked and tdTOMATO intensity was scored at selected time points. EOM and TA MuSCs displayed distinct profiles of lineage progression, where TA myogenic cells initiated reporter gene expression earlier than EOM cells (**Figure 1E**). The percentage of *Myog*-expressing cells sharply increased in the TA from 5% at 80h post-plating to 40% at 96h. Significantly, only ∼7% of EOM cells were tdTOMATO^+^ by 96h *in vitro* (**Figure 1F**).

Since *Myog* expression is followed by cell-cycle withdrawal and terminal differentiation (Andrés and Walsh, 1996; Benavente-Diaz et al., 2021; Guo et al., 1995), a delay in its expression could allow for sustained proliferation of EOM MuSCs. We performed RT-qPCR at 48h post-plating and assessed the expression of key myogenic transcription factors and the cell cycle inhibitor *p21*, which regulates cell cycle exit during myoblast differentiation (Guo et al., 1995; Zhang et al., 1999). In agreement with the live imaging data, *Myog* mRNA was significantly upregulated in TA activated MuSCs compared to EOM, concomitant with an increase in *p21* mRNA (**Figure 1G**). No significant differences were found in *Pax7* nor *Myod* expression (**Figure 1G**). Moreover, EOM progenitors displayed reduced levels of phosphorylated P38 (**Suppl Fig1C,D**). As p38α/β MAPKs play a central role in promoting MEF2 transcriptional activity and initiating the differentiation program (Rugowska et al., 2021), this data suggests that EOM progenitors are less prone to myogenic commitment. Therefore, our results suggest that a delay in the differentiation of EOM MuSCs might contribute to the higher number of myogenic cells produced compared to TA cells.

### Activation of EOM MuSCs include a larger progenitor pool and distinct activation signatures

To gain insights into the potential mechanisms that regulate the phenotypic differences and cell fate decisions between EOM and TA MuSCs, we performed scRNA-seq of *in vitro* activated MuSCs using the 10x Chromium platform. We first isolated EOM and TA MuSCs by FACS from *Tg:Pax7-nGFP* mice and cultured them *in vitro* for 4 days. Cell cultures were subsequently trypsinized, cell-sorted and processed for scRNA-seq (**Figure 2A**). Unsupervised clustering divided cells into 2 clusters per sample (**Figure 2B**) that were annotated as progenitors (Prog) or differentiating (Diff) based on the expression of known myogenic makers, such as *Pax7/Myf5* and *Myod/Myog* (**Figure 2B, C**). Notably, expression of Myod was lower in EOM progenitors (**Figure 2B**) and lower protein levels were detected at the whole population level in vitro (**Suppl Figure 2A**).

**Figure 2.**
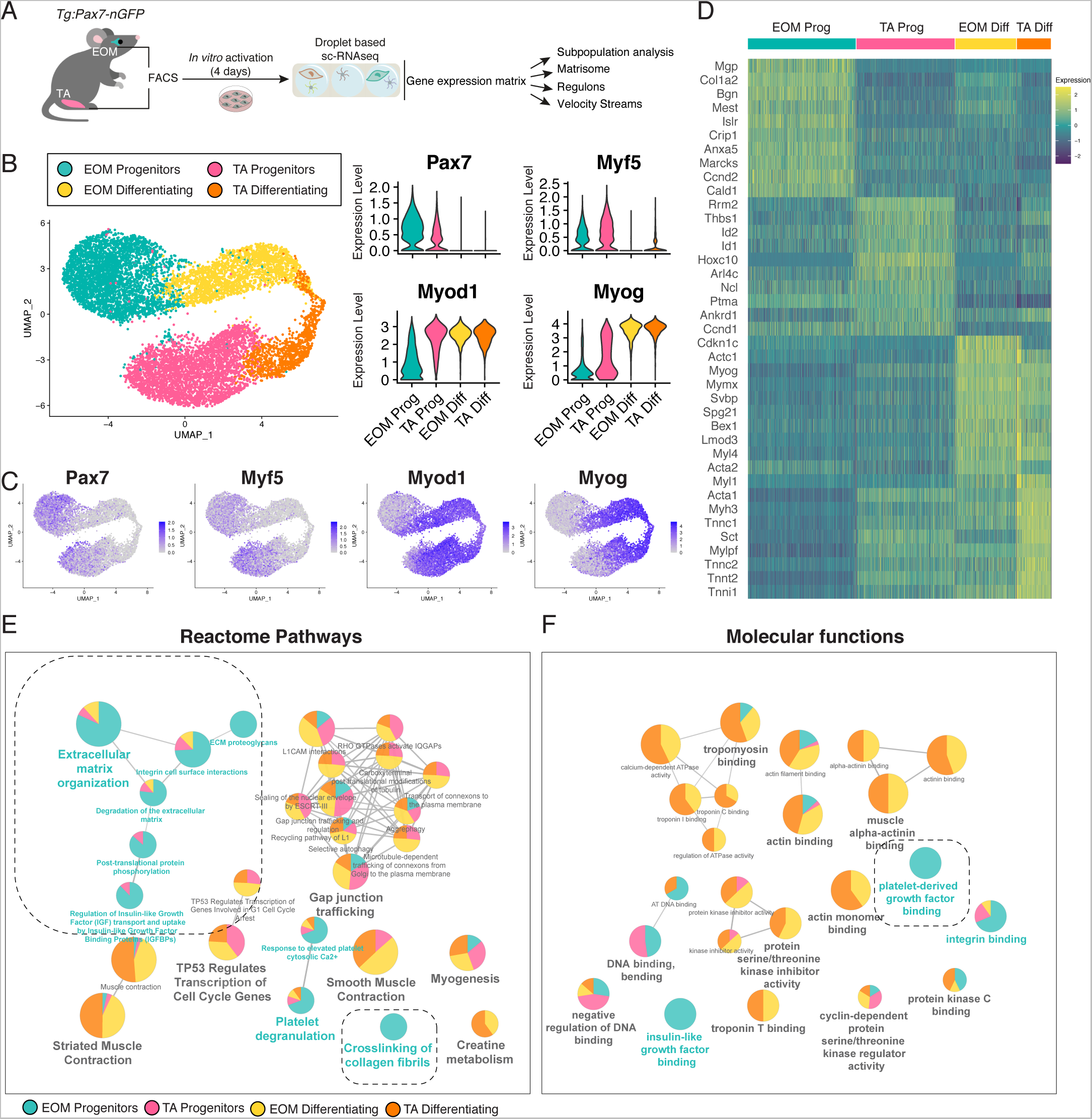
Single cell transcriptome signatures of activated EOM and TA MuSCs. A. Experimental scheme of sc-RNAseq pipeline for in vitro activated EOM and TA MuSCs. B. UMAP visualization of EOM progenitors, EOM differentiating, TA progenitors, and TA differentiating cell clusters (left). Violin plots of the expression of myogenic markers *Pax7* (progenitor), *Myod1* (committed) and *Myog* (differentiating) in each cluster (right). C. Expression plots of myogenic markers for progenitors (*Myf5*, *Pax7*) and committed/differentiating cells (*Myod*, *Myog*). D. Heatmap representing top differentially expressed genes in each cluster and expression levels across all cells. E-F. Reactome pathway (E) and GO Molecular function (F) network analysis on top 100 DEGs of each cluster. Pie charts represent relative contribution of each cluster to this ontology term. EOM-specific selected terms are highlighted.

Analysis of differentially expressed genes (DEGs) across these clusters showed that each cell state had a distinct transcriptional pattern (**Figure 2D, Suppl Table 1**). As expected, TA progenitors expressed a defined Hox-signature (Evano et al., 2020) and inhibitors of differentiation (Id1, Id2, (Jen et al., 1992; Kumar et al., 2009)). Instead, EOM progenitors displayed markers that had not been previously described in activated MuSC datasets such as *Mgp*, *Bgn*, *Col1A2* and *Acta2* (smooth muscle actin). EOM and TA differentiating cells shared part of their signature (*Myog, Mymx, Myl4*). However, TA progenitors shared some markers with the TA differentiating cells (*Acta1, Mylpf, Tnnt2*), suggesting that Prog and Diff TA clusters are closer at a transcriptional level than their EOM counterparts (**Figure 2D**).

To get insights into these properties and the prospective functional heterogeneity associated with each transcriptomic signature we performed reactome pathway analysis of the DEGs upon in vitro activation (**Figure 2E, Suppl Figure 2B, Suppl Table 2**). Multiple pathways involved in ECM organisation were characteristic of EOM progenitors including “ECM proteoglycans”, “crosslinking of collagen fibrils” and “integrin cell surface interactions”. On the other hand, EOM and TA differentiating cells were both enriched in pathways involved in striated muscle contraction and creatinine metabolism, which are associated with maturation of skeletal muscle. Also, as suggested by the DEG heatmap, TA progenitors shared common pathways with the differentiating clusters. Analysing the molecular function of the DEGs revealed that EOM progenitors were also specifically enriched in Pdgfrβ, a receptor for platelet-derived growth factor, and proteins related to insulin like growth factor binding and integrin signaling (**Figure 2F, Suppl Figure 2C, Suppl Table 3**). Thus, these analyses uncovered a non-canonical signature in the EOM progenitor fraction upon activation.

### EOM and TA MuSCs retain distinct molecular signatures after several days in culture

We then set out to determine to what extent the EOM progenitor fraction that was capable of self-renewal *in vitro*, relates to the quiescent MuSCs present *in vivo*. Thus, we assessed the profiles of quiescent MuSCs by scRNA-seq of from EOM and TA immediately after isolation by FACS (**Figure 3A-B**). Despite the homogeneous expression of MuSC markers *Myf5* and *Pax7* in both populations, *Myod* expression was restricted to the TA MuSCs (**Figure 3C-D**), suggesting an earlier MuSC activation in response to the isolation procedure (Brink et al., 2017; Machado et al., 2017; Velthoven et al., 2017). Differentiation markers like *Myog* were absent in both populations (**Figure 3C-D**). Visualization of the top 25 most variably expressed genes documented distinct transcriptional programs of EOM and TA quiescent clusters (**Figure 3E, Suppl Table 4**). Moreover, microarray analysis of adult quiescent *Tg:Pax7-nGFP* MuSCs revealed that the EOM quiescent signature was also different from that of MuSCs from other cranial-mesoderm derived muscles such as the esophagus and masseter (**Suppl Figure 3A**). Interestingly, we found several conserved genes between the quiescent and activated sc-RNAseq states at each anatomical location, while others were unique to the quiescent or activated cell states (**Figure 3F-I**). In the TA, *Lbx1, Vgll2* and *Hox* genes were found as common signature throughout cell states (**Figure 3G,I, Suppl Figure 3B, C**). *Lbx1* is a homeobox transcription factor required for the migration of myogenic progenitor cells to the limbs (Brohmann et al., 2000; Gross et al., 2000) and *Vgll2* (previously named Vito-1) is a key cofactor of the myogenic differentiation program (Günther et al., 2004; Maeda et al., 2002). All three genes had been already identified in the limb both at the MuSC levels and in entire muscles in the adult (Evano et al., 2020; Honda et al., 2017; Terry et al., 2018). *Pcdh7*, an integral membrane protein belonging to cadherin superfamily that regulates intercellular adhesion (Wang et al., 2020; Yoshida, 2003) is unique to TA quiescent state and the microtubule protein *Tubb3* (Duly et al., 2022) is a marker of the TA activated state (Figure 3I).

**Figure 3.**
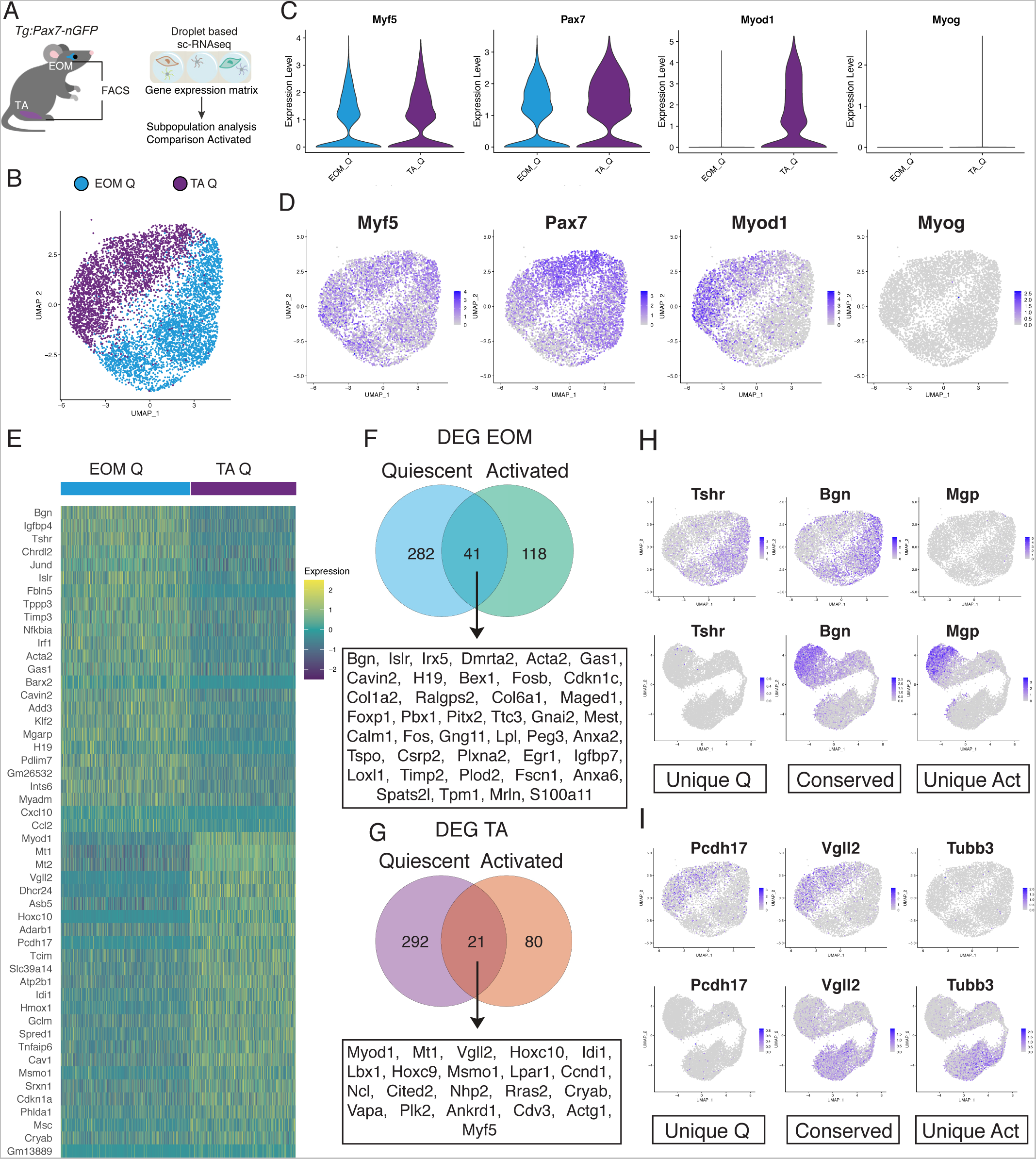
Molecular signature of EOM and TA quiescent MuSCs. A. Experimental scheme of sc-RNAseq pipeline for quiescent EOM and TA MuSCs. B. UMAP visualization of EOM and TA quiescent clusters. C-D. Violin plot (C) and expression plot (D) of *Myf5*, *Pax7*, *Myod* and *Myog*. E. Heatmap representing top differentially expressed genes in EOM and TA quiescent clusters and their expression levels across all cells. F-G. Venn diagrams of the overlapping differentially expressed genes between quiescent and global in vitro activated datasets for EOM (F) and TA (G). H. Expression plots of selected EOM markers exclusive to the quiescence state (Unique Q), common to quiescent and activated states (conserved) and exclusive to activation (Unique Act) from the analysis in F. I. Expression plots of selected TA markers exclusive to the quiescence state (Unique Q), common to quiescent and activated states (conserved) and exclusive to activation (Unique Act) from the analysis in G.

EOM genes unique to the quiescent state included *Tshr*, encoding the thyroid-stimulating hormone receptor, which has been previously identified in bulk RNAseq as “EOM-resistant” kept upon engraftment into the limb (Evano et al., 2020), and shown to control senescence in MuSCs of DMD rats (Taglietti et al., 2023). Matrix Gla protein (Mgp), which is a critical regulator of angiogenesis in multiple organs (Kida and Yamaguchi, 2022), was identified as exclusively upregulated in EOM cells upon activation (Figure 3H). Amongst the conserved EOM genes across cell states we identified *Pitx2*, a well-known major upstream regulator of EOM development (Gage et al., 1999), *Fos* gene family members (Fos and Fosb), with *Fos* being recently identified in a subset of limb MuSCs with enhanced regenerative capacity (Almada et al., 2021), and *Igfbp7*, a specific marker of quiescent MuSCs that is downregulated upon activation (Fukada et al., 2007) but also reported to be upregulated in MuSCs upon exercise (Chen et al., 2020) (**Figure 3F, Suppl Figure 3B, C**). Additionally, the EOM common signature included several ECM components and regulators (e.g. *Bgn*, *Loxl1*, *Col1a2*, *Col6a1*), TFs associated with fibrosis and connective tissue development (e.g. *Foxp1* (Grimaldi et al., 2022; Shao and Wei, 2018), *Egr1* (Havis and Duprez, 2020)) and *Acta2* (alpha smooth muscle actin), a marker of smooth muscle, fibroadipogenic progenitors (Joe et al., 2010; Uezumi et al., 2010) and smooth muscle-mesenchymal cells (SMMCs, (Giordani et al., 2019)) (**Figure 3F,H**). Finally, other top markers for EOM MuSCs identified in the quiescent microarray including *Foxc1,* which is involved in ocular development (Smith et al., 2000) and reported to regulate the balance between myogenic and vascular lineages within somites (Lagha 2009,Mayeuf-Louchart 2016), *Eya2, Islr* and *Igfbp4*, were also enriched in the activated EOM sc-RNAseq dataset (**Suppl Figure 3B**). Altogether, our analysis revealed a closer molecular overlap for EOM MuSCs across cell states, including several TFs and ECM markers that continued to be distinct from TA MuSCs.

### EOM MuSC activation involves extensive ECM remodelling and a mesenchymal-like signature

Given the well-established role of ECM synthesis and remodelling on MuSC proliferation and self-renewal (Baghdadi et al., 2018; Rayagiri et al., 2018; Tierney et al., 2016; Yin et al., 2013), we sought to further characterise the EOM and TA signatures using the Matrisome database (Matrisome DB) (Naba et al., 2015), which compiles *in silico* and experimental data on ECM constituents. We first identified the components of the Matrisome DB present in our single-cell dataset and created a global score for ECM component expression (**Figure 4A**) (see Material and Methods). Significantly, the EOM progenitor cluster had the highest matrisome score, while no major differences where observed within the other clusters (**Figure 4A**). Next, we identified the matrisome components that were differentially expressed in each cluster and found the EOM progenitor cluster also expressed the highest number of genes for each matrisome category when compared to the other clusters (**Figure 4B**).

**Figure 4.**
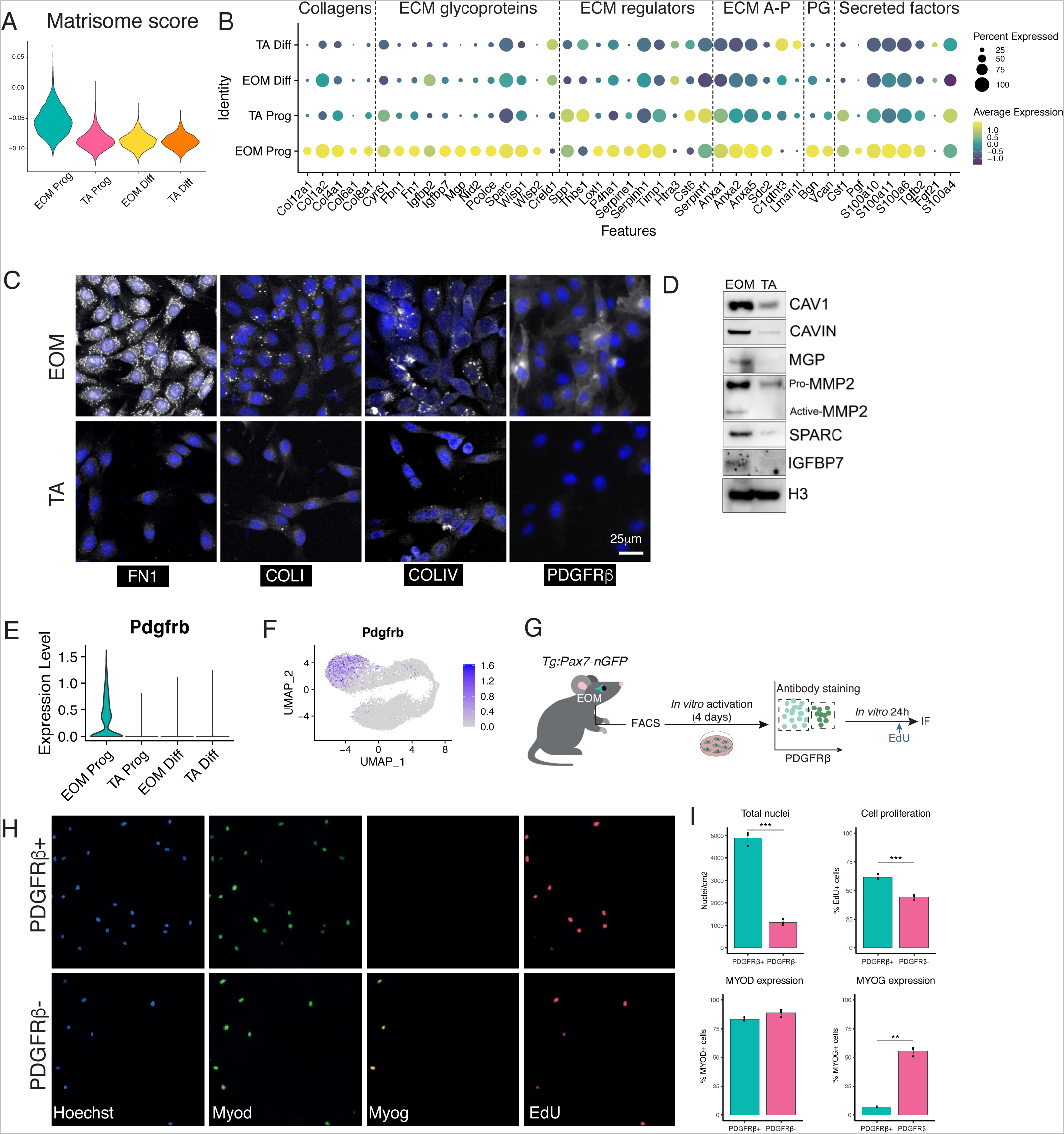
EOM MuSC activation is accompanied by ECM remodelling. A. Violin plots of matrisome scores for each cluster of the activated sc-RNAseq dataset. B. Dot plot visualisation of differentially expressed matrisome components of the activated sc-RNAseq dataset. ECM, extracellular matrix; AP, affiliated proteins; PG, proteoglycans. C. Immunofluorescence for Fibronectin (FN1), Collagen I (COLI), Collagen IV (COLIV) and Pdgfrα. D. Western blot showing expression of CAV1, CAVIN1, MGP, MMP2, SPARC, IGFBP7 and H3 for normalization. Cells coming from n=3 mice pooled per lane. E. Violin plot of *Pdgfrα* expression on in vitro activated sc-RNAseq dataset. F. UMAP of *Pdgfrα* expression on in vitro activated sc-RNAseq dataset. G. Scheme of isolation and culture of MuSCs isolated from adult *Tg:Pax7-nGFP* mice. EOM cells were resorted based on their PDGFRβ expression, plated for 24h and pulsed with EdU before fixation for immunostaining. H. Representative images of PDGFRβ positive and negative EOM MuSC fractions stained for MYOD, MYOGENIN and EdU as described in J. I. Bar plots of quantification of number of cells/cm^2^ and percentage of EdU^+^, MYOD^+^ and MYOGENIN^+^ cells. n=3 mice, >100 cells counted. Scale bar 50 μm. Two-tailed unpaired Student’s t-test; ** p-value<0.01, *** p-value<0.005.

We then validated some of the matrisome candidate genes at the protein level in activated MuSCs. EOM and TA MuSCs were isolated by FACS, cultured *in vitro* for 4 days and protein expression was assessed by immunofluorescence or Western blot (**Figure 4C-D**). Immunostaining for Fibronectin (FN1) showed a marked difference in fluorescence intensity between EOM and TA cells, confirming that EOM cells express higher levels of this protein (**Figure 4C**). Given that FN1 secretion by MuSCs promotes MuSC expansion in a cell autonomous manner (Bentzinger et al., 2013), the distinct expression of *Fn1* might contribute to maintenance of the progenitor population during EOM MuSC activation. EOM progenitors also had higher levels of Collagen I (COLI) (**Figure 4C**), a major component of the fibrotic ECM (Dulauroy et al., 2012) shown to suppress differentiation of C2C12 cells and display mutually exclusive expression with Myogenin (Alexakis et al., 2007). Similarly, EOM progenitors expressed higher levels of Collagen IV (COLIV) (**Figure 4C**), which was reported to be secreted by MuSCs as well as myoblasts and fibroblasts in culture (Baghdadi et al., 2018; Kühl et al., 1984). In addition, PDGFRβ, a tyrosine-kinase receptor commonly expressed by mesenchymal cells and pericytes (Hellström et al., 1999; Levéen et al., 1994; Soriano, 1994), was also differentially expressed in EOM progenitors by immunofluorescence (**Figure 4C**). Interestingly, Pdgfrβ and Acta2 were described as a markers of Smooth Muscle-Mesenchymal Cells (SMMCs), a novel Itga7+ Vcam-Pdgfrb+ Acta2+ cell subpopulation present in adult muscle that exhibits myogenic potential and promotes MuSC engraftment following transplantation (Giordani et al., 2019).

Western blot analysis confirmed that EOM progenitors produce higher levels of Caveolin1 (CAV1) and Cavin1 (**Figure 4D**), which are co-expressed in caveolae and shown to be downregulated upon differentiation of rhabdomyosarcoma cells (Faggi et al., 2015). CAV1 was also shown to be a marker of quiescent and Pax7+ activated mouse MuSCs (Gnocchi et al., 2009) whereas in human, the CAV1+ MuSC subpopulation is associated with ECM organization, expression of quiescent markers and display increased engraftment after transplantation (Barruet et al., 2020). SPARC, MGP and IGFBP7 were also found to be upregulated in EOM activated MuSCs by Western blot (**Figure 4D**). The activity of these proteins seems to be context dependent and they can promote or suppress proliferation in different cell types (Ahmad et al., 2017; Artico et al., 2021; Cho et al., 2000; Jing et al., 2019; Kuronuma et al., 2020; Li et al., 2020; Melouane et al., 2018; Said et al., 2013). Given the large number of matrisome genes differentially expressed in EOM progenitors, we also examined the expression of MMP2, a matrix remodelling protein (Gonçalves et al., 2022). Western Blot analysis revealed an enrichment in the active form of MMP2 in EOM progenitors whereas the TA progenitors displayed an enrichment of the pro-MMP2 or latent form (**Figure 4D)**. This result is in agreement with the observation that activation of MMP2 in vitro and in vivo increases the proportion and mobility of Pax7+ cells (Mu et al., 2010).

In parallel, we generated various gene signature scores to assess to what extent activated EOM MuSCs resemble other cells described in skeletal muscle displaying mesenchymal features (**Suppl Fig 4, Suppl Table 5**). As expected, EOM progenitors displayed a higher score for smooth muscle and mesenchymal cells (SMMCs, (Giordani et al., 2019)), FAPs (Oprescu et al., 2020), myotendinous junction B myonuclei (Kim et al., 2020), Twist2+ population(Liu et al., 2017), fetal MuSCs (Tierney et al., 2016), developing limb connective tissues (Lima et al., 2021) and the skeletal muscle mesenchyme identified in human fetal limb (Xi et al., 2020). Instead, TA progenitors and the respective EOM and TA differentiated fractions displayed a higher score for myogenic commitment and differentiation (**Suppl Fig 4**).

Finally, we focused on Pdgfrα for further analysis as it is a component of the matrisome (**Figure 4B**) and a gene identified on the EOM molecular functions (**Suppl Figure 2C**) differentially expressed in EOM progenitors (**Figure 4E,F**). Pdgfrβ is a tyrosine-kinase receptor commonly expressed by pericytes (Hellström et al., 1999; Soriano, 1994) and it is involved in multiple signalling pathways that regulate cell migration, proliferation, and differentiation (Heldin and Westermark, 1999). We isolated this subpopulation by FACS from in vitro activated EOM cells using a PDGFRβ antibody. The flow cytometry analysis corroborated the scRNA-seq and immunofluorescence data (**Figure 4C**), where PDGFRβ was enriched in EOM samples (**Suppl Figure 4B,C**). To distinguish the functional properties of PDGFRβ+ and PDGFRβ-EOM cells, we sorted both subpopulations from EOM cultures, which represent respectively 83 and 17% of total nuclei at D4 and cultured them in growth promoting conditions for an additional 24h (**Figure 4G-I**). Cells were treated with EdU for 2h prior to fixation to monitor cell proliferation. PDGFRβ+ cells had a significantly higher proliferative capacity as measured by the total nuclear output and incorporation of EdU compared to PDGFRβ-cells (**Figure 4H,I**). Although the percentage of MYOD-expressing cells was not overtly different between the two populations, MYOG was expressed by over 50% of PDGFRβ-cells, whilst being detected in only 7% of PDGFRβ+ cells (**Figure 4H,I**). Therefore, the PDGFRβ+ myoblast subpopulation is characterised by a higher proliferative potential and decreased differentiation status.

Altogether, this analysis showed that EOM progenitors have an unusual transcriptome profile, express a wide range of ECM-related factors and harbour a mesenchymal signature that may endow them with a higher remodelling capacity of their immediate niche and promotion of an undifferentiated state.

### EOM transcriptome profile is associated with a unique transcription factor network

To identify the underlying programs of myoblast heterogeneity between the two muscle groups, we inferred single cell regulatory networks using pySCENIC (Aibar et al., 2017; Sande et al., 2020). SCENIC uses co-expression patterns and transcription factor binding motifs to expose ’regulons" (transcription factors and their putative targets). This pipeline generates a new matrix comprising the activity of each regulon, in each cell (**Figure 5A, Suppl Table 6**). As expected, regulons associated with myoblast differentiation such as *Myod*, *Myog* and *Mef2* family were found to be specifically active in differentiated cells of both EOM and TA (**Figure 5B**). Of note, the top 5 regulons of TA progenitors were also found to be active in EOM progenitors, whereas the top 5 regulons of EOM progenitors were unique to this cluster (**Figure 5B**). EOM progenitors were characterised by a unique signature of regulons involved in connective tissue/ECM remodelling including *Egr1* (Havis and Duprez, 2020) and Creb3l1, a downstream effector of Thr signaling (García et al., 2017) that plays a critical role during bone development and activates extracellular matrix genes such as Col1a1 and Fn1 (Sampieri et al., 2019). Top regulons of EOM progenitors are also involved in cell proliferation (*Foxc1* (Yang et al., 2017), *Sox4* (Moreno, 2019), *Fos* (Almada et al., 2021), *Klf6* (Dionyssiou et al., 2013), *Ebf1* (Györy et al., 2012)), commitment into endothelial and smooth muscle fates (*Foxc1* (Han et al., 2017; Mayeuf-Louchart et al., 2016; Whitesell et al., 2019; Yang et al., 2017)) as well as differentiation into mesenchymal lineages such as pericytes, adipocytes and chondrocytes (Ebf1, (El-Magd et al., 2021; Jimenez et al., 2007; Pagani et al., 2021). Notably, some of these regulons were also active in the EOM during development and implicated in non-myogenic cell fates from bipotent myogenic cells (Grimaldi et al., 2022). Interestingly also, for some of these EOM progenitor regulons, the genes themselves were already enriched in quiescent EOM MuSCs and their expression was upregulated upon activation (**Suppl Fig 5A, B**).

**Figure 5.**
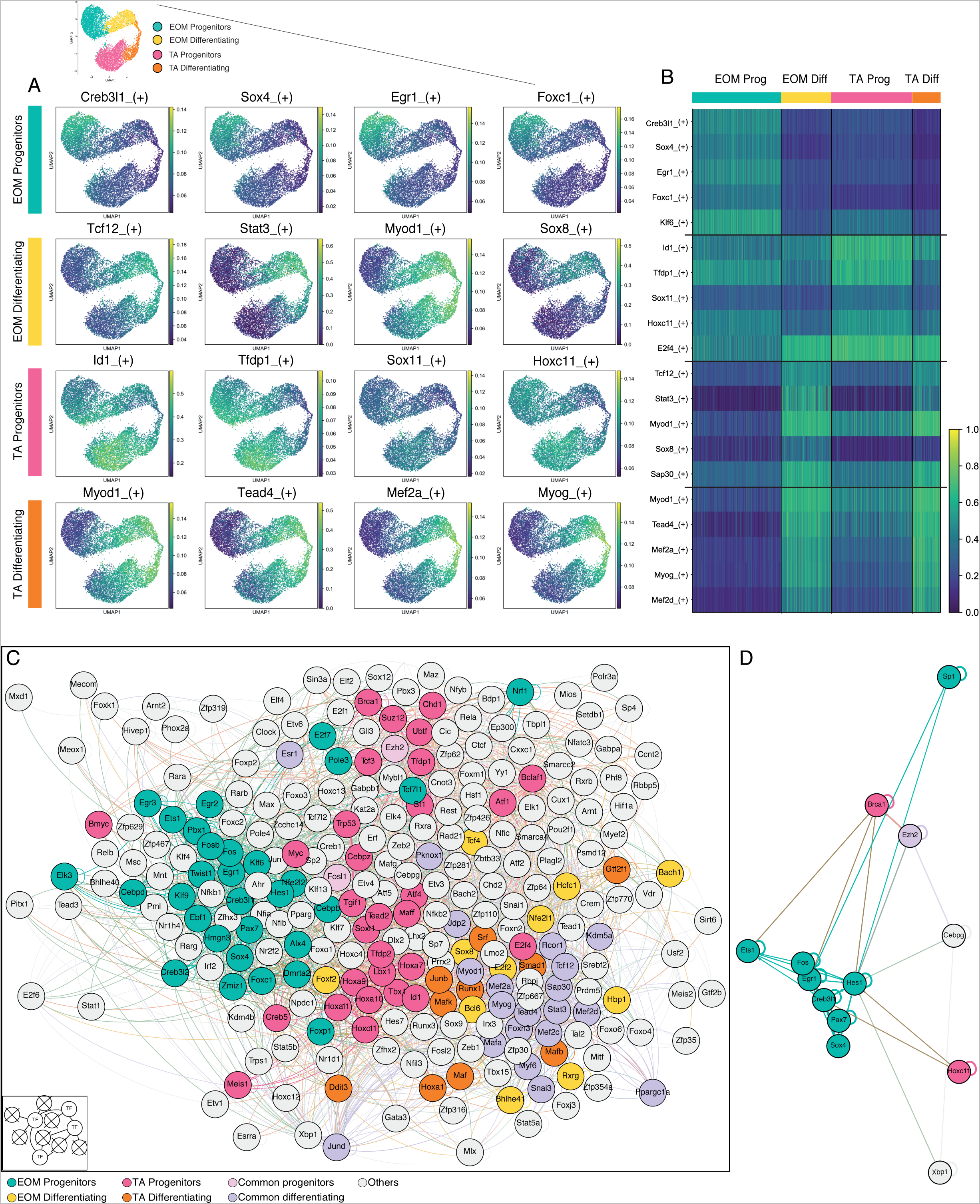
Distinct gene regulatory networks underlie EOM and TA activation dynamics. A. Top 4 regulon activity for each cluster of in vitro activated sc-RNAseq dataset overlaid onto UMAP representation. B. Heatmap of Top 7 regulons in each cluster of the activated sc-RNAseq dataset with activity level in each cell C. Transcription factor network highlighting top regulons of each cluster as well as common regulons (35 regulons maximum) of activated sc-RNAseq dataset. Proximity of the nodes in the network indicate a higher number of shared edges, highlighting core modules. D. Filtered network showing Hes1 direct regulators (nodes with 1 level ancestry).

To visualise the potential interactions between these regulons, we built a network restricted to transcription factors. To do so, target genes that were not at the head of regulons themselves were removed (Grimaldi et al., 2022) (see Methods) (**Figure 5C).** This results in a layout that visually represents the relationships between the nodes, where each node (circle) is an active transcription factor and each edge (distance between nodes) is an inferred regulation between 2 transcription factors. When placed in a force-directed environment (see Methods), these nodes aggregated based on the number of shared edges. Thus, we were able to highlight associated and co-regulating transcription factor modules. Strikingly, the transcription factors of the most specific regulons of each cluster preferentially organised as tightly related modules (**Figure 5C**). In agreement with our previous analyses, the known co-regulating transcription factors found in differentiated cells (*Myod, Myog, Mef2a, Mef2c, Myf6*) formed a tight module in this visualisation (**Fig 5C**, common differentiating category). Interestingly, the TA muscle progenitor module was composed of genes that were reported to be required for limb embryonic development (*Hox* genes, *Lbx1*) (Gross et al., 2000; Pineault and Wellik, 2014; Swinehart et al., 2013). Notably, *Hoxa11*, is a determinant of limb axial identity during embryonic development (Zakany and Duboule, 2007), and HoxA and HoxC clusters were found as signatures of adult TA MuSCs (Evano et al., 2020; Yoshioka et al., 2021); **Suppl Fig5B**).

Within the EOM progenitor module, the network included *Foxc1, Egr1*, *Creb3l1*, *Dmrta2*, *Sox4*, *Fos* and *Egr1* transcription factors together with *Pax7* and *Hes1* which are known to support the maintenance of quiescence (Baghdadi et al., 2018; Mourikis et al., 2012; Olguin and Olwin, 2004; Relaix et al., 2005). In this context though, the network seems to maintain a proliferative progenitor state in activated EOM MuSCs cells as assessed by the higher protein levels of PAX7, CCND1 (Cyclin D1), together with other EOM regulon TFs (FOXC1, EBF1 and CREB3L1) (**Suppl Fig5C, D**). The vicinity of Hes1 to the EOM regulatory module prompted us to explore its direct upstream regulators (one level of ancestry in the network) (**Figure 5D**). Strikingly, 8 potential activators (including *Hes1* itself) were part of the EOM module. By contrast, only 2 upstream regulators were part of the TA regulatory module (**Figure 5D**). Of note, *Fos* and *Egr1* have also been reported as part of a stress signature following tissue dissociation (Machado et al., 2021). Yet, in our dataset, the expression of these genes (i.e. the StressIndex) was correlated with the expression of *Pax7* and anti-correlated with the expression of *Myod* (**Suppl Fig5E,F**). Moreover, when we applied a regression of the StressIndex or removed these genes from the count matrix, the general aspect of the data did not change (**Suppl Fig5G)**, pointing at a role of these genes in the EOM progenitor maintenance independently of the stress response.

### EOM features are present during the growth phase in vivo and retained upon passages in vitro

From the analysis above, it appears clear that EOM MuSCs possess an unusual transcriptomic state that may repress myogenic commitment and maintain a more "stem-like" state. We then wondered whether these EOM features are cell-wired and persist upon passage in vitro. Cells were isolated from *Tg:Pax7-nGFP;Myog^ntdTom^*mice and analyzed upon in vitro activation or upon one or two passages (**Figure 6A, Suppl Fig 6A, B**). While the total cell number was reduced with passages for both EOM and TA MuSCs (**Suppl Fig 6A, B**), the normalized cellular output was consistently higher for the EOM and correlated with a lower tdT/GFP ratio (**Figure 6B, C**). Real-time qPCR analysis revealed that the expression of Pax7 and Hey1, a bHLH transcription factor that is required in a cell-autonomous manner for maintenance of MuSCs (Noguchi et al., 2019), EOM specific regulon TFs (Foxc1, Sox4, Ebf1, Creb3l1) and genes identified by the matrisome or molecular functions (Bgn, Sparc, Igfbp2, Igfbp7, Pdgfrb) were retained or even increased in EOM cultures after several passages compared to activated EOM MuSCs cultured only for 4 days (**Figure 6D**). These results were confirmed at the protein level by Western blot analysis (**Figure 6E, F**) where the specific EOM signature described above, that seems to maintain a more proliferative state in activated EOM MuSCs cells, persists even after several passages in vitro. Indeed, EOM MuSCs cells retained higher protein levels of PAX7, EOM regulon TFs (**Figure 6E**) as well as EOM matrisome or molecular functional genes (**Figure 6F**) compared to TA MuSCs cells at same passage. These results show that EOM MuSCs cells retain a cell autonomous non-canonical signature that is hard-wired even after extended cell culture.

**Figure 6.**
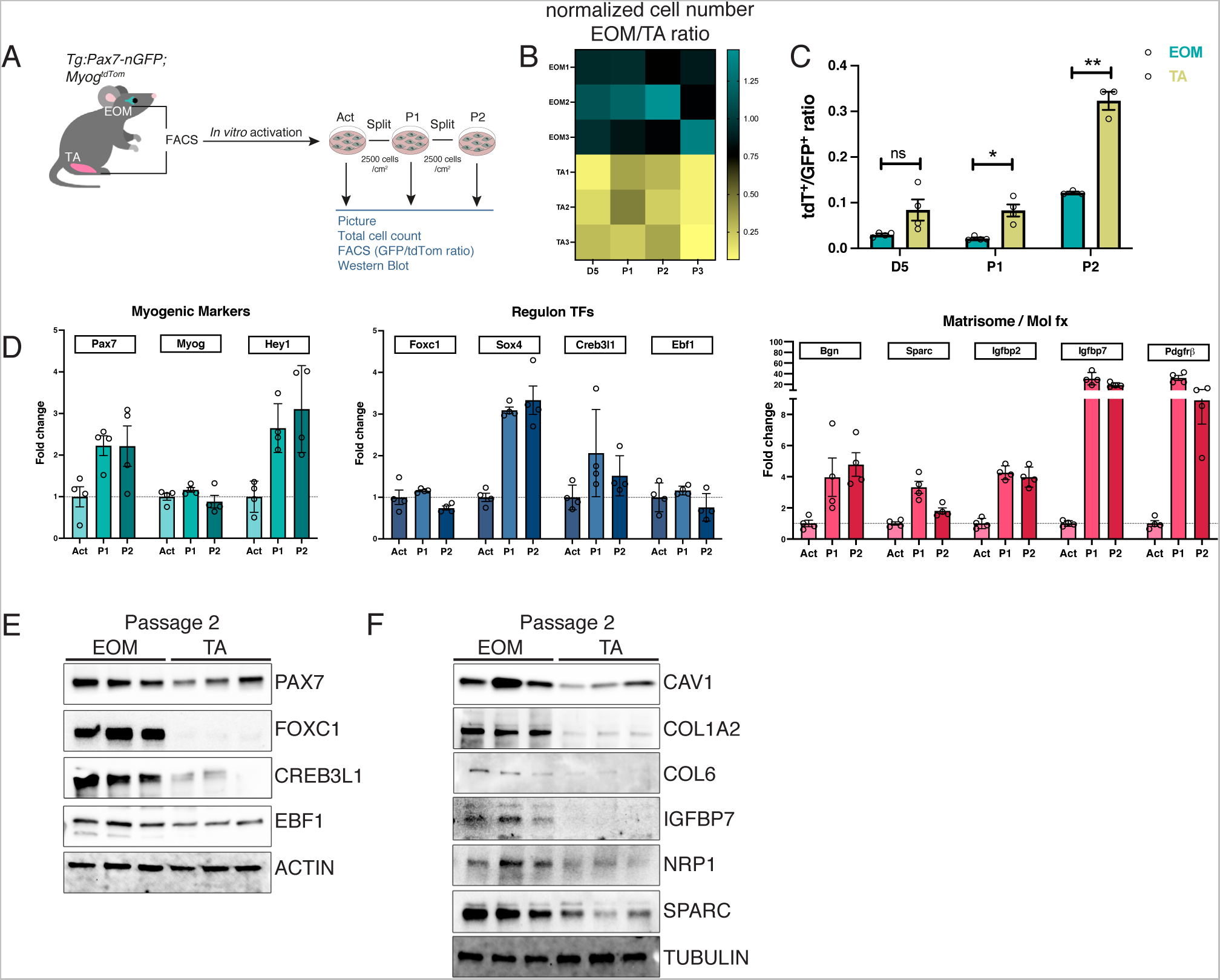
EOM properties are retained despite several passages in vitro. A. Scheme of isolation and passages of EOM and TA MuSCs. Cells were isolated by FACS based on GFP fluorescence from *Tg:Pax7-nGFP;Myog^ntdTom^* and cultured for 3 days (Act). The entire well was passaged (P1, P2) every 3 days. Following each passage, a fraction of cells was analysed by FACS to assess GFP and tdTom ratios. B. EOM/TA normalized cell number ratios. Total cell numbers per well were counted upon trypsinization at day 3 (D3), days 5 (D5) and at passages (P)1 and P2. EOM/TA normalized cell number ratio was generated for each time point analysed (n=3 mice). For normalization, the ratio between cell numbers counted per well and averaged EOM cell number in all wells at each time point was calculated. C. FACS analysis of *Tg:Pax7-nGFP;Myog^ntdTom^* EOM and TA MuSCs upon 5 days in culture (D5), one or two passages (P1, P2) (n≥3 mice). Graph represents ratio of TdT^+^/GFP^+^ cells for each time point. Two-tailed unpaired Student’s t-test; *p-value<0.05, ** p-value<0.01. D. qRT-PCR at D3 (Act, Activated), P1, P2 on whole populations present in dish for key myogenic markers, regulon TFs and Matrisome/Molecular function genes identified in EOM progenitors (n=4 mice). E. Western blot of total P2 population of EOM and TA MuSCs (n=3 mice; one mouse per lane). Note higher expression of Pax7 and EOM progenitor specific TF regulons (FOXC1, CREB3L1, EBF1) identified in activated EOM MuSCs by sc-RNAseq. ACTIN was used for normalization of protein loading. F. Western blot of the total P2 population of EOM and TA MuSCs (n=3 mice; one mouse per lane) showing higher expression of matrisome genes (CAV1, COL1A2, COL6, IGFBP7, NRP1, SPARC) identified in activated EOM MuSCs by sc-RNAseq. ACTIN was used for normalization of protein loading.

Next, we asked whether EOM progenitor features are already present in activated fetal development and postnatal stages, where extensive muscle growth and MuSC expansion occurs, or if these features were acquired in adulthood upon reactivation from the quiescent state. To address these possibilities, we first isolated EOM and TA MuSCs from E18.5 and P21 *Tg:Pax7-nGFP* mice and performed qRT-PCR for Myogenin after 48h in culture (**Suppl Figure 7A**). As observed for adult MuSCs (**Figure 1D**), EOM cultures were less differentiated. However, as GFP protein persists during myogenic commitment (Sambasivan et al., 2009) and this can introduce a bias in the initial populations, we isolated the GFP+/tdT-fractions from *Tg:Pax7-nGFP;Myog^ntdTom^* mice at P7 by FACS (**Figure 7A, Suppl Figure 7B**) and performed RT-qPCR. Similarly, to the *in vitro* sc-RNAseq data, significantly lower levels of *Myogenin* were detected in EOM GFP+/tdT-cells (**Figure 7B**). Finally, we confirmed higher transcript levels for the EOM specific regulon TFs (*Foxc1*, *Ebf1*, *Sox4*, *Creb3l1*) and matrisome or molecular functional components identified by sc-RNAseq of activated MuSCs *in vitro* (**Figure 7B**). Altogether, our results indicate that EOM MuSCs repress myogenic commitment and maintain a more "stem-like" state upon activation, and this property is conserved from development to adulthood.

**Figure 7.**
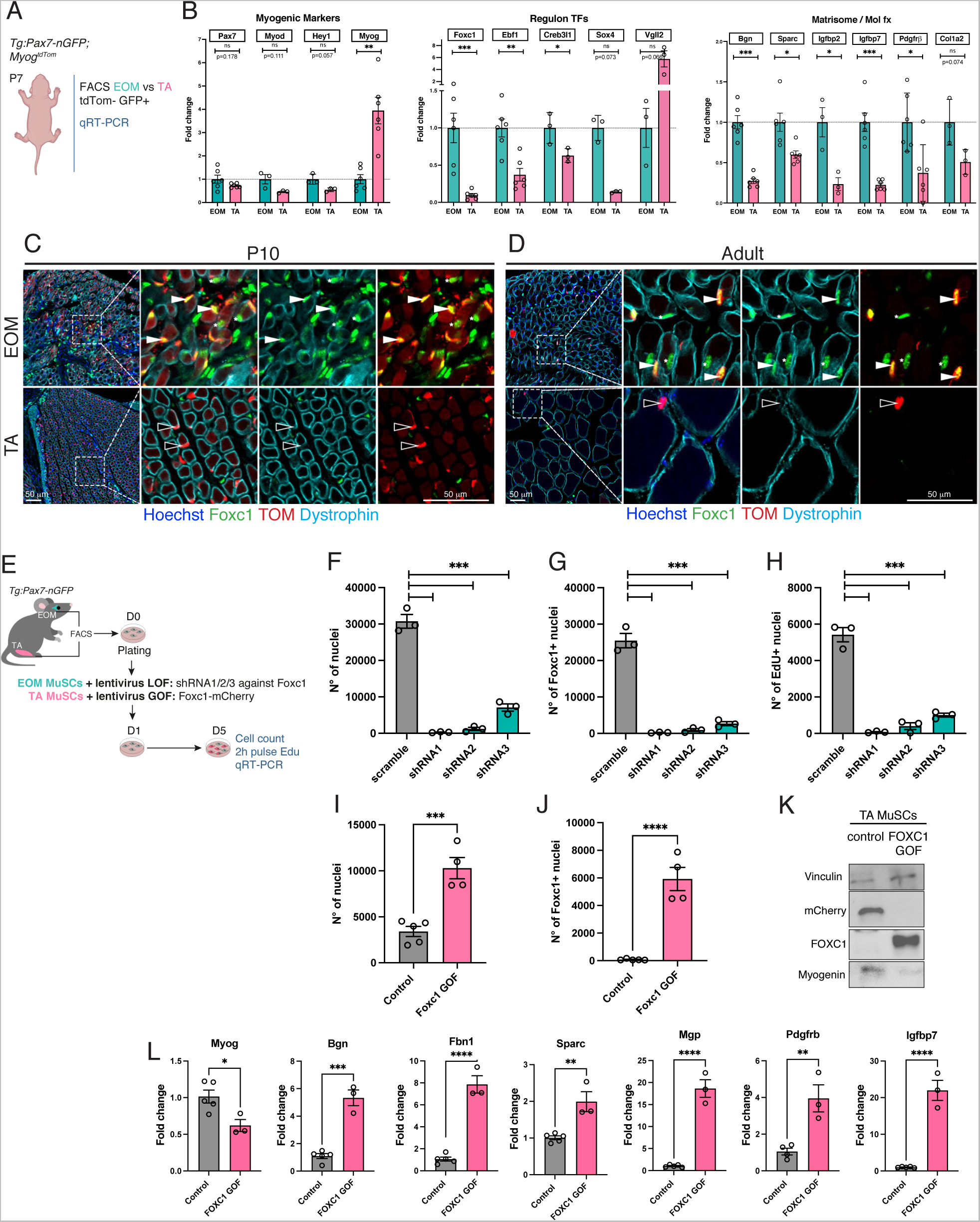
EOM activated features are present during postnatal growth and can be modulated by Foxc1. A. Scheme of isolation of EOM and TA MuSCs from *Tg:Pax7-nGFP;Myog^ntdTom^* from postnatal day (P) 7 mice. B. qRT-PCR for key myogenic markers, regulon TF and Matrisome/Molecular function genes identified in EOM progenitors by scRNAseq analysis. GFP^+^/TdT^-^ Cells were isolated from *Tg:Pax7-nGFP;Myog^ntdTom^* EOM and TA MuSCs at postnatal day (P) 7 (n=3-6). C-D. Immunostaining for FOXC1, tdTOMATO (TOM) and Dystrophin on cryosections from EOM and TA muscles isolated from P10 (C) and adult (D) *Pax7^CreERT2^:R26^tdTom^* mice (n=2 per stage). White arrowheads point to PAX7+FOXC1+ cells. Black arrowheads point to PAX7+FOXC1-cells. E. Scheme of lentivirus transduction for loss of function (LOF) of EOM and gain of function (GOF) of TA MuSCs. Cells from *Tg:Pax7-nGFP* mice were isolated by FACs, plated and transduced the next day (MOI 100). Cells were analysed at D5. F-H. Bar plots of quantification of number nuclei, FOXC1+ nuclei and EdU+ nuclei of EOM MuSCs transduced with scramble or shRNA against Foxc1 containing viruses. I-J. Bar plots of quantification of number nuclei and FOXC1+ nuclei of TA MuSCs transduced with control or Foxc1 expressing viruses. F-L, Two-tailed unpaired Student’s t-test; *** p-value<0.005, **** p-value<0.0001. K. Western blot of TA MuSCs transduced with control or Foxc1 expressing viruses for mCherry, Foxc1 and Myogenin. Vinculin was used for normalization of protein loading. L. qRT-PCR of TA MuSCs transduced with control or Foxc1 expressing viruses for Myogenin and key ECM proteins and regulators (Creb3l1) identified in EOM progenitors.

### Foxc1 marks the EOM MuSC lineage and plays a role in progenitor cell maintenance

We then focused on Foxc1 for further analysis as it is one of the top regulons and DEG of the activated sc-RNAseq dataset (**Figure 5 A,B, Suppl Figure 3B, Suppl Figure 5B, D**) and expressed continuously in the EOM lineage. Notably also, bulk RNAseq from adult muscles (Terry et al., 2018) showed that Foxc1 expression is higher in entire EOMs than in any of the other 10 muscle groups analyzed (**Suppl Fig 7C**). Finally, immunostaining on tissue sections of *Pax7^CreERT2^:R26^tdTOM^*mice displayed colocalization of FOXC1 with tdTOMATO in EOM MuSCs and expression of FOXC1 in EOM myonuclei at P10 and in adulthood but not in the TA (**Figure 7C, D**). Thus, differences in Foxc1 expression between the EOM and TA seem to arise during development and are kept upon activation, and throughout the myogenic lineage.

As differences in *Myogenin* expression at the mRNA level are already evident at Day2 upon in vitro activation (**Figure 1G**), we decided to use this early timepoint to functionally validate a potential role of Foxc1 in the maintenance of the progenitor state. Thus, we silenced *Foxc1* via a siRNA approach in EOM activated MuSCs (**Suppl Figure 7D-F**). While the total cell number and percentage of Pax7 cells was not changed 2 days after Foxc1 silencing, immunostaining for FOXC1 showed silencing of 82% of the cells at the protein level (**Suppl Figure 7E**) and RT-qPCR showed a 64% reduction at the transcript level (**Suppl Figure 7F**). Moreover, a 2.7 fold increase in *Myogenin* expression was detected by qPCR following silencing of *Foxc1* as early as Day2 (**Suppl Figure 7F**). To look at later timepoints, we transduced EOM activated MuSCs with lentiviruses expressing different short-hairpin RNAs against Foxc1 (**Figure 7E**). Immunofluorescence and EdU uptake at D5 revealed an efficient depletion of Foxc1 protein, and concomitant severe reduction in the total cell number, and number of EdU+ cells (**Figure 7F-H**). We then overexpressed Foxc1 in activated TA MuSCs to assess whether overexpression of a single factor could confer at least a subset of EOM features to TA cells (**Figure 7E, I-L**). This approach resulted in a 3-fold increase in the total cell number (**Figure 7I**), number of FOXC1+ cells (**Figure 7J**) and reduction of Myogenin protein levels (**Figure 7K,L**). As *Foxc1* direct targets (**Suppl Table 7**) include matrisome components (e.g. *Sparc*, *Pdgfrβ*, *Fbn1*, **Figure 4B**) and other EOM specific regulon TFs (e.g. *Ebf1*, *Creb3l1*, *Egr1*, **Figure 5B**), we assessed expression of these genes upon *Foxc1* overexpression (**Figure 7L**). Many of the matrisome genes assessed were upregulated in this context.

Altogether, our findings support the notion that *Foxc1* is a key TF for maintenance of EOM MuSCs in a progenitor like state and matrix deposition. Overexpression of this factor in TA cells seems to be sufficient to recapitulate part of the EOM in vitro activated phenotype.

### Transcription dynamics expose EOM and TA disparities in progenitor-state maintenance

We first attempted to correlate the number of regulons responsible for most of the matrisome gene expression differences between EOM and TA upon activation (**Figure 8A-C, Suppl Table 8**). We observed that activated EOM MuSCs consistently regulated a higher number of matrisome genes than TA cells independently of the number of regulons considered (**Figure 8A**). However, the ratio of the number of regulations of matrisome genes between EOM and TA activated MuSCs showed that the difference was maximal when considering the top 5 regulons, *Alx4*, *Dmrta2*, *Foxc1*, *Zmiz1* and *Fos* (**Figure 8B, C**). Notably, Dmrta2, Foxc1 and Fos were also active regulons in quiescence, with slight disparities between EOM and TA (**Figure 8D, Suppl Table 9**).

**Figure 8.**
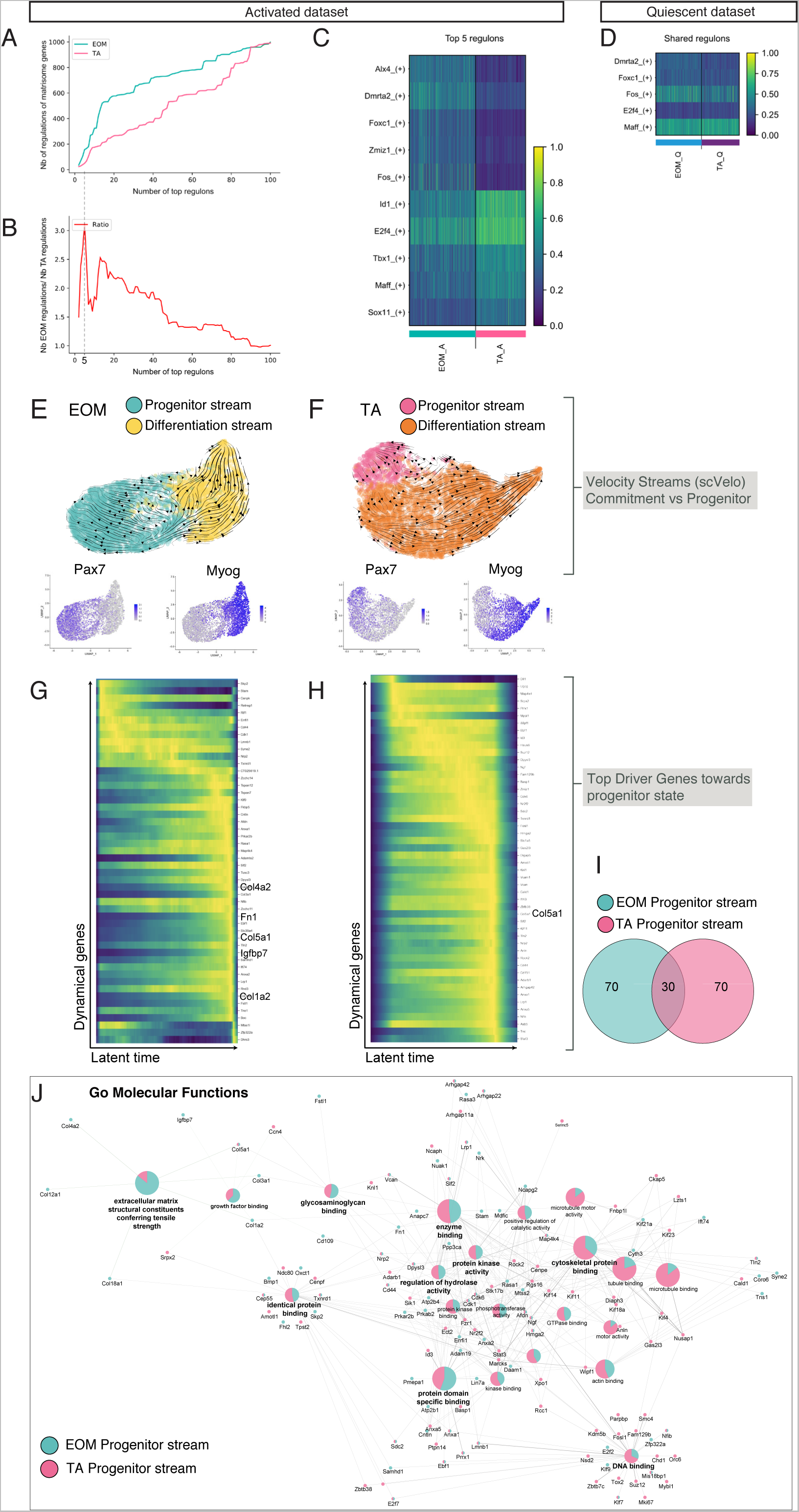
RNA velocity reveals distinct population kinetics and potential key regulators of progenitor maintenance. A. Number of regulatory links between regulons and matrisome genes depending on the number of top regulons in EOM and TA in vitro activated sc-RNAseq datasets. B. Ratio of number of regulations of matrisome genes between EOM and TA MuSCs on in vitro activated sc-RNAseq dataset. Note difference is maximal for top 5 first regulons. C. Heatmap of top regulons in global in vitro activated sc-RNAseq dataset with activity level in each cell. D. Heatmap of same top 5 regulons in quiescent sc-RNAseq dataset. Note activities in quiescence is similar between EOM and TA, but seem to diverge during activation. E. EOM Velocity streams overlaid onto a UMAP representation, along with expression patterns of *Myog* and *Pax7* on in vitro activated sc-RNAseq dataset. F. TA Velocity streams overlaid onto a UMAP representation, along with expression patterns of *Myog* and *Pax7* on in vitro activated sc-RNAseq dataset. G. Heatmap of driver genes expression, obtained from EOM progenitor velocity stream. H. Heatmap of driver genes expression, obtained from TA progenitor velocity stream. I. Venn diagram of top 100 drivers for EOM and TA progenitor streams. J. GO Molecular function ontology network and associated driver genes exposing unique and common pathways dynamically regulated during EOM and TA progenitor maintenance.

Second, we set out to determine whether matrix genes underlie the transition towards progenitors and committed cells during activation. To do so, we inferred RNA velocity using scVelo (Bergen et al., 2020). This method computes local changes in the relative amount of unspliced and spliced variants, which depend on the rates of transcription, degradation and splicing (Manno et al., 2018). ScVelo identifies candidate "driver genes", i.e. the most transcriptionally dynamic genes in a given cluster that are responsible for most of the inferred RNA velocity, and thus underlie its transitional trajectory. We applied the scVelo pipeline to EOM and TA datasets independently, and examined the directional trajectories underlying commitment and progenitor maintenance (**Figure 8E,F**). Two distinct velocity streams stood out in both datasets, towards differentiation (*Myog*^High^) and towards a progenitor-state (*Pax7*^High^). Strikingly, a larger fraction of EOM cells appeared to transition towards a progenitor state compared with the TA (**Figure 8E,F**). Conversely, most cells in the TA appeared to be directed towards differentiation, consistent with an overlapping transcriptomic profile with differentiating cells, shown as top DEGs (**Figure 2D**) and regulatory networks (**Figure 5C**). These trajectories did not appear to be specifically correlated with cell cycle phases (**Suppl Figure 8 A,B**), which was shown to influence transcriptomic data in some cases (McDavid et al., 2016). Hence, the velocity streams observed are most likely to reflect transitions between distinct cell states instead of the cell cycle progression of a homogeneous cell state.

Using scvelo built-in functions, we extracted the top driver genes underlying the velocity towards the progenitor state in both datasets (**Figure 8G-I, Suppl Table 10, 11**). Out of the top 100 driver genes, 30 were common to both datasets, including *ColV*, which plays a critical role in maintenance of quiescence (Baghdadi et al., 2018) (**Figure 8G,H**). We assessed the GO molecular functions associated with these driver genes to identify the unique and shared pathways employed by both datasets during myogenic progenitor maintenance (**Figure 8J**). In agreement with our previous results, this transition was characterised by the active upregulation of ECM components specifically in the EOM.

In summary, our results establish a new gene regulatory network (GRN) permitting an enhanced expansion capacity and delayed commitment to differentiation of EOM MuSCs upon activation. This regulatory module is hardwired to EOM identity and relies on the active maintenance of progenitor MuSC features.

## DISCUSSION

One of the fundamental, yet unanswered questions in stem cell and regenerative biology regards the mechanisms that allow an orchestrated balance between proliferation, differentiation and self-renewal. An unusual feature of skeletal muscle stem cells is their reliance on distinct GRNs in different anatomical locations. How these pathways confer the unique functional properties of each stem cell population in cranial and limb/limb muscles remains poorly understood. Indeed, most studies on adult and developmental myogenesis have focused on trunk and limb muscles and only a handful of transcription factors and signalling pathways have been identified as hallmarks of specific muscle groups. In this study we performed scRNA-seq accompanied by experimental validations to characterise the GRN underlying the functional heterogeneity of MuSCs derived from different muscles, taking as EOM and TA as archetype muscles for high and low performing stem cell populations.

### EOM MuSCs are more refractory to *in vitro* differentiation

Although subsets of cranial derived MuSCs were previously described to be more proliferative and to have a higher engraftment capacity than trunk MuSCs (Ono et al., 2010; Randolph et al., 2015; Stuelsatz et al., 2015), the mechanisms responsible for these phenotypic differences remained elusive. As these features depend on the balance between proliferation and differentiation, one possibility is that different upstream regulators define the pace at which MuSCs progress in the lineage in diverse anatomical positions.

Here, by monitoring *Myog* expression, we showed that EOM MuSCs have a lower propensity to differentiate following activation and thus persist as a proliferative population. As foetal and early postnatal EOM MuSCs are also refractory to differentiation, we propose that this property might be hardwired to some extent by unique GRNs that are retained throughout development and adulthood. Notably, foetal MuSCs were shown to be more resistant to myogenic progression upon in vitro expansion and contribute more efficiently upon transplantion than the adult counterparts (Sakai et al., 2013; Tierney et al., 2016). In addition, a recent study showed that limb MuSCs cells isolated at birth displayed prolonged expansion rate and a delay in differentiation and fusion compared with those isolated at later stages of postnatal growth (P7 and P15) and adulthood (Gattazzo et al., 2020). Therefore, is tempting to speculate that EOM MuSCs retain features of the respective foetal and neonatal precursor cells.

EOM MuSCs are strong candidates for cell-based therapies given their robust engraftment capacity in vivo (Stuelsatz et al., 2015). So far, major obstacles in the clinic are the large cell numbers required for transplantation and the fact that ex vivo amplification of trunk MuSCs leads to a drastic decline in regenerative potential due to commitment to differentiation (Briggs and Morgan, 2013; Ikemoto et al., 2007). Substantial advances have been made by modifying the MuSC culture conditions (Charville et al., 2015; L’honoré et al., 2018), or by the use of teratoma-derived skeletal myogenic progenitors (Xie et al., 2021). As such, the identification of factors that regulate cell fate decisions in distinct MuSC populations serves as a resource for advancing knowledge in the context of regenerative medicine.

### EOM progenitors exhibit a mesenchymal genetic signature upon activation

Recent scRNA-seq analyses have provided some insights into the transcriptional landscape regulating MuSC quiescence, activation and self-renewal in somite-derived muscles (Dell’Orso et al., 2019; Hernando-Herraez et al., 2019; Machado et al., 2021; Micheli et al., 2020; Yartseva et al., 2020). To characterise these processes in cranial MuSCs, we performed a comparative scRNA-seq analysis of activated EOM and TA MuSCs. Distinct transcriptional profiles divided the myoblast pool into two subpopulations: those that resembled progenitors and those that were differentiating. The EOM progenitor population was characterised by a higher *Pax7* levels, and thus reminiscent of *in vitro* reserve cells (Laumonier et al., 2017; Yoshida et al., 1998; Zammit et al., 2004) and ontology analysis of its DEGs revealed a surprising enrichment in ECM organisation processes and *Pdgf* signalling.

By surveying bulk RNA sequencing and single-cell profiling data of trunk skeletal muscle stem cells, a recent study highlighted a dynamic profile of Pdgf ligands and receptors during myogenesis (Contreras et al., 2021). In addition, treatment with NOTCH and PDGFRβ ligands (DLL4 and PDGF-BB respectively) was shown to enhance migration, expression of stem cell markers and perivascular-like features in MuSCs (Gerli et al., 2019) and embryonic myoblasts (Cappellari et al., 2013). Given that PDGFRβ and several Notch pathway components are co-expressed in EOM progenitors, and at higher levels than the TA, we speculate that cross-talk between these pathways could take place in this subpopulation.

In addition to *Pdgfrβ,* which is a well-known pericyte marker (Lindahl et al., 1997), EOM progenitor cells express *Acta2* which is involved in the contractile apparatus of smooth muscle and also used as a SMMCs (Pdgfrb+ Itga7+ Vcam- Pdgfrb+ Acta2+) and FAPs (Giordani et al., 2019; Joe et al., 2010; Uezumi et al., 2010) marker. However, while the EOM progenitor fraction expressed *Fbn1* and *Loxl1*, which are markers of a new subtype of FAPs identified by scRNA-seq of whole muscle (Rubenstein et al., 2020), they were negative for canonical FAPs and pericyte markers such as *Pdgfrα* (Joe et al., 2010; Uezumi et al., 2010) and *Cspg4* (*Ng2*) (Birbrair et al., 2013), respectively. In addition, the EOM progenitor fraction also related more to fetal MuSCs than the TA counterparts in terms of enrichment in matrisome components such as Fn1, Fbn1, Vcam and collagens (Tierney et al., 2016). It is well established that cell autonomous ECM deposition and remodelling are key instructive cues governing cell fate decisions in MuSCs (Baghdadi et al., 2018; Lukjanenko et al., 2016; Mashinchian et al., 2018; Tierney et al., 2016). Thus, we hypothesize that EOM progenitors might exploit these features to preserve self-renewal potential and mimic their in vivo stem cell niche even after in vitro culture. In the EOM, this plasticity has already been observed in vivo, where EOM myogenic progenitors transition towards non-myogenic cell fates in areas deprived of neural crest cells during mouse embryogenesis (Grimaldi et al., 2022). Additionally, a resident subpopulation expressing reduced levels of canonical myogenic markers but increased levels of mesenchymal markers has been detected in embryonic and foetal human limb muscle (Xi et al., 2020). This plasticity seems to be present also *in vitro*, where directed myogenic differentiation of hiPSCs generates heterogeneous cell types including both myogenic and non-myogenic cells (Xi et al., 2020).

### A unique network of transcription factors maintains EOM progenitors

Analysis of single cell regulatory networks (SCENIC) revealed a unique set of regulons whose activity was restricted to EOM progenitors. Of note, the majority of the identified head regulon genes in EOM progenitors are not typical myogenic TFs. One of the most active regulons in this cluster was *Foxc1*, which was described as a pro-mitogenic factor in the cancer field (Yang et al., 2017) and as driver of endothelial/smooth muscle fate alone (Whitesell et al., 2019) or in combination with Foxc2 (Mayeuf-Louchart et al., 2016). Another top regulon was *Egr1*, which was shown to promote the expression of many ECM-related genes in different tissues (Gaut et al., 2016; Guerquin et al., 2013; Milet et al., 2017; Wu et al., 2017). Interestingly, *Foxc1* and *Ebf1* were found as active transcription factors underlying the transition of myogenic towards non-myogenic cell fates in the EOM anlage during embryonic development (Grimaldi et al., 2022). Both of these transcription factors were found in EOM progenitors in this study, suggesting that they might be continuously active from development to adulthood specifically in the EOM. Interestingly, expression of Foxc1 by hair follicle stem cells activates quiescent gene networks upon activation (Wang et al., 2016). As *Foxc1* is expressed both by quiescent and activated EOM MuSCs, this gene might reinforce a progenitor/stem-cell identity throughout the lineage. Notably, *Pitx2* appears as a target of *Foxc1* in our analysis (Suppl Table 7) and its overexpression promotes MuSC proliferation in vitro and enhances the regenerative potential of dystrophic MuSCs in vivo (Vallejo et al., 2018). Other non-canonical myogenic TF identified in EOM progenitors are the Kruppel-like factor family members *Klf4*, *Klf6*, and *Klf9* with *Klf6* identified as top regulons. This gene family participates in the development and homeostasis of numerous tissues (McConnell and Yang, 2010), and *Klf4* is a pioneer factor well-known for its reprogramming capabilities (Schmidt and Plath, 2012)

When assessing the interactions between transcription factors, the EOM Progenitor regulon TFs clustered together in a regulatory module including *Pax7* and the Notch target *Hes1*, supporting our hypothesis that these cells are in a more stem/progenitor state. This module also included the transcription factor Fos/AP-1, whose expression in a subset of freshly isolated trunk MuSCs is related to enhanced regenerative properties such as rapid cell cycle entry, higher clonal capacity and more efficient engraftment (Almada et al., 2021). Further, by means of an inducible FOS expression system, it was reported that elevated FOS activity in recently activated muscle progenitor cells perturbs cellular differentiation by altering the 3D chromosome organization near critical pro-myogenic genes (Barutcu et al., 2022). Altogether, it is tempting to speculate that a common regulatory module allows the establishment and maintenance of a MuSC subpopulation in the EOMs that displays mesenchymal features, partially overlaps with the quiescent state, and is refractory to differentiation.

To investigate the maintenance of this stem state during EOM activation we used RNA velocity to infer the different directional trajectories and their driver genes. The proportion of cells transitioning to a more progenitor state was higher in the EOM when compared to TA, and this progenitor stream was driven by a specific gene set. Amongst these genes there was an enrichment of ECM-related factors, underscoring its active role in the maintenance of a less differentiated state. Interestingly, we identified *Igfbp7* (Insulin growth factor binding protein 7), as unique EOM driver gene, which has been found to be a specific marker of quiescent MuSCs (Fukada et al., 2007). Taken together, our results highlight an active maintenance of a progenitor pool that partially overlaps with the quiescent state during EOM MuSC activation, possibly through the steady induction of inhibitory differentiation signalling cues and promotion of ECM remodelling.

## Conclusion

Using *in vitro* and *in silico* approaches, we propose a model where the outperformance of EOM MuSCs depends on the expression of a tightly associated module of transcription factors regulating a characteristic and distinct pattern of ECM-remodelling factors, cell receptors and growth factor binding proteins. These components define the pace at which EOM MuSCs progress through the myogenic lineage and maintenance of a stem-like population. As such, our study lays the groundwork for elucidating the mechanisms of selective sparing of muscle groups in dystrophic disease by providing information on a unique core GRN within this muscle group. In addition, our study could serve as basis for the optimization of in vitro protocols that would exploit cranial MuSC properties in the context of cell-based muscle therapies.

## Materials and methods

### Animal care

Animals were handled according to national and European Community guidelines and an ethics committee of the Institut Pasteur (CETEA) in France approved protocols. Except when indicated otherwise, males and females of 2-4 months were used. *Tg:Pax7-nGFP* (Sambasivan et al., 2009), *Myog^ntdTom^*(Benavente-Diaz et al., 2021), *Pax7^CreERT2^* (Mathew et al., 2011) and *R26^tdTom^* (Madisen et al., 2009) mouse lines were maintained in a C57Bl/6JRj background. To induce recombination of *Pax7^CreERT2^:R26^tdTom^* mice a 20 mg/ml stock solution of Tamoxifen was prepared in 5% ethanol and 95% sunflower seed oil by thorough resuspension with rocking at 4 °C. For adult mice, 2 mg of tamoxifen (Sigma #T5648) were administered by gavage during 5 consecutive days and animals sacrificed 5 days later. To induce recombination of pups, the Tamoxifen stock solution was diluted to 15mg/ml with sunflower seed oil and 0.15 mg were administered daily by subcutaneous injection between P4 and P6 daily. Pups were sacrificed at P10 by decapitation and adult mice by cervical dislocation.

### MuSC isolation by FACS

Muscles were dissected and minced in ice-cold DMEM. Samples were then incubated in DMEM (Fisher Scientific, 11594446), 0.08% Collagenase D (Sigma, 11088882001), 0.2% Trypsin (ThermoFisher, 15090) and 10 µg/mL of DNAse I (Sigma, 11284932) for 25 minutes at 37°C under gentle agitation for 5 rounds of digestion. After each round, samples were allowed to precipitate for 5 minutes, the supernatant was collected on 4mL of Foetal Bovine Serum (FBS) on ice and fresh digestion buffer was added to the remaining muscle pellet. The collected supernatants were centrifuged for 15 minutes at 550 g at 4°C, resuspended in DMEM 2% FBS and filtered through a 40µm strainer (Corning, 352235) before cell sorting.

Cells were isolated based on size, granularity and GFP fluorescence using an Aria III (BD Biosciences) sorter. Cells were collected directly in MuSC growth media (DMEM: F12 (1:1, GIBCO), 20% FBS (ThermoFisher, #10270), 2% Ultroser (Pall, 15950-017), 1% Penicillin/Streptomycin (GIBCO, 15140-122).

For myoblast isolation based on PDGFRβ expression, dishes were washed twice with PBS prior to trypsinization. Cells were subsequently incubated with α-PDGFRβ antibody in HBSS (Fisher Scientific, 11560616) media with 2% FBS for 30 minutes on ice. After two washes with HBSS 2% FBS, cells were incubated with streptavidin-coupled BV786 and incubated on ice for 25 minutes. After one wash with HBSS 2% FBS, the cell suspension filtered through a 40µm strainer (Corning, 352235) before cell sorting. Cells were isolated based on size, granulosity and GFP or BV786 fluorescence on using an Aria III (BD Biosciences) machine.

### Muscle stem cell culture, treatment and transfection

Cells isolated by FACS were plated onto Matrigel® (Corning, 354248) coated dishes and cultured in growth media at 3% O2, 5% CO2, 37°C for the indicated time. To assess proliferation, cells were pulsed with 10^-6^ M of EdU (ThermoFisher, C10640) in cell culture media for 2h prior to fixation.

For loss of function experiments, freshly isolated satellite cells from Tg:Pax7-nGFP mice were transfected in suspension immediately after FACS with the ON-TARGET plus SMARTpool against FOXC1 (Dharmacon, L-047399-01-0005) or Scramble (Dharmacon, ON-TARGETplus Non-targeting Control siRNA, D0018100205) at 200 nM final concentration using Lipofectamine 3000 (ThermoFisher, L3000001) in Opti-MEM (Gibco) as described by the manufacturer. Briefly, a pre-mix of siRNA/Optimem (1.5ul /20ul) and Lipofectamine3000/Optimem (0.3ul / 20ul) were incubated separately 5 min RT, mixed at 1:1 ratio and incubated 15 min more at RT. 2.10^4^ cells in 40ul of Optimem were incubated with an equal volume of the transfection mix for 2h at 3% O2, 5% CO2, 37°C in Eppendorf tubes whose caps had had been punctured with a needle to allow gas exchange. Two hours after transfection, three volumes of fresh growth medium were added and cells were plated at 10k/cm2 in Matrigel coated wells containing growth media. Two days upon transfection wells were processed for immunostaining or RNA collected with Tryzol as above.

### Lentivirus transduction for gain and loss of function experiments

Control (mCherry), Foxc1-mCherry and Foxc1-shRNA containing vectors were made by VectorBuilder (https://en.vectorbuilder.com/). Freshly isolated satellite cells from *Tg:Pax7-nGFP* mice were plated in matrigel coated wells, cultured overnight and transduced at an MOI (multiplicity of infection) of 100 in 45μl (for 96 well plates) or 125μl (for 48 well plates) of MuSC media containing 5 μg/ml of polybrene. After 4h incubation at 37°C, cells were washed three times with MuSC and cultures for 4 more days prior to fixation for immunostaining, or protein or RNA collection.

### RT-qPCR

RNA from in vitro myoblast cultures was extracted using a Trizol-based kit (Zymo Research, R2061) and reverse transcribed using SuperScriptIII (Invitrogen, 18080093). RT-qPCR to assess for mRNA relative expression performed with SYBR green master mix (Roche, 04913914001) in an Applied biosciences machine. Data analysis was performed using the 2-ΔΔCT method (Livak and Schmittgen, 2001) and mRNA expression was normalized with *Rpl13*.

### Immunofluorescence

For immunostaining of ECM markers, cells were fixed in warm (37°C) 3% paraformaldehyde (PFA, Electron Microscopy Sciences) 2% sucrose. Upon incubation in the fixative for 15min at RT, cells were washed three times with PBS and incubated with NH4Cl 100mM in PBS to quench aldehyde groups. Cells were washed three times in PBS, permeabilized in 0.5% Triton X-100 (Merck, T8787) for 5 min RT, washed again three times in PBS and blocked with 10% goat serum (GIBCO). Cells were incubated with the indicated primary antibodies overnight at 4°C in PBS with 2% goat serum. Upon PBS washes, cells were incubated with secondary antibodies and Hoechst (ThermoFisher, H1399) for 45 min at RT. EdU was detected after antibody staining if needed according to manufacturer instructions (ThermoFisher, C10640).

Following immunostaining, 96-well plates (Dutscher Dominique, 655090) were imaged on an Opera Phenix high-content microscope (Perkin Elmer) using a 20x or 40x objective. Quantification of acquired images was performed using Harmony (Perkin Elmer) or Columbus software, using an automated pipeline. First, nuclei were detected based on Hoechst signal. Number of nuclei was assessed and mean and maximum intensity were automatically quantified on the nuclear region for nuclear markers. For acquisition of ECM markers either the Opera Phenix high-content microscope (Perkin Elmer) or Zeiss LSM800 microscope with ZEN software were used.

Immunostaining on tissue sections was performed as (Comai et al., 2020). Immunostaining for FOXC1 on tissue sections required an amplification step using Alexa Fluor 488 Tyramide SuperBoost Kit (IgG rabbit, ThermoFisher, B40943).

### Western Blot

Cells in culture were lysed in RIPA buffer (150mM NaCl, 50mM Tris pH8, 1mM EDTA, 1% Triton X100 (Sigma, T8787), 0.5% sodium deoxycholate, 0.1% SDS supplemented with 1X proteases (Sigma, S8820) and phosphatases (Roche, 4906845001) inhibitors) and kept on ice for 30 minutes. The homogenate was then centrifuged for 10 min at 3000g, 4°C. The protein concentration was determined by Bradford assay (Bio-Rad, 5000205) using a BSA standard curve. The protein absorbance was measured at 595 nm by using a microplate reader Infinite M2000 (Tecan). Equal amounts of protein were reconstituted in 4x Laemmli Sample Buffer (Bio-Rad, 1610747) and heated at 95°C for 5 min. 5 μg of protein extracts were run on a 4%–15% polyacrylamide gels (Mini Protein TGX Stain-Free gels, Bio-Rad) and transferred to PVDF (Bio-Rad, 1704156) membrane with Trans-Blot® Turbo™ Transfer system (Bio-Rad). The membrane was blocked with 5% milk or BSA (PAN-Biotech, P06-1391100) in Tris-Buffer Saline 0.2% Tween (Sigma, P9416) (TBS-T) for 1h at room temperature and probed with specific primary antibodies overnight at 4°C. After three washes in TBS-T, the membrane was incubated with HRP or fluorophore-conjugated secondary antibodies and revealed by chemiluminescence (Clarity Max Bio-Rad, 1705062) or fluorescence (Bio-Rad, Chemidoc MP) and detected using the ChemiDoc® Gel Imaging System. Densitometric analysis of the immunoblots was performed using Image Lab Software v.6.1.0 (Bio-Rad).

### Time lapse microscopy

MuSC were plated on a microscopy culture chamber (IBIDI, 80826) and cultured in growth media. The plate was incubated at 37°C, 5% CO_2_, and 3% O_2_ (Zeiss, Pecon). A Zeiss Observer.Z1 connected to a Plan-Apochromat 20x/0.8 M27 objective and Hamamatsu Orca Flash 4 camera piloted with Zen (Zeiss) was used.

### Image analysis

Cell tracking was performed using the Manual Tracking feature of the TrackMate plug-in (Tinevez et al., 2017) in Fiji (Schindelin et al., 2012). Fiji and Harmony (Perkin Elmer) were used for image analysis. Figures were assembled in Adobe Photoshop, Illustrator and InDesign (Adobe Systems).

### Data analysis and statistics

Data analysis, statistics and visualisations were performed using Prism (Graphpad Software) or using R (Team, 2014) and the package ggplot2 (Wickham, 2009). For comparison between two groups, two tailed unpaired Student’s t-tests were performed to calculate p values and to determine statistically significant differences (see Figure Legends).

### scRNAseq data generation

MuSCs were isolated on BD FACSAria™ III based on GFP fluorescence and cell viability from *Tg:Pax7-nGFP mice* (Sambasivan et al., 2009). Quiescent MuSCs were manually counted using a hemocytometer and immediately processed for scRNA-seq. For activated samples, MuSCs were cultured *in vitro* as described above for four days. Activated MuSCs were subsequently trypsinized and washed in DMEM/F12 2% FBS. Live cells were re-sorted, manually counted using a hemocytometer and processed for scRNA-seq.

Prior to scRNAseq, RNA integrity was assessed using Agilent Bioanalyzer 2100 to validate the isolation protocol (RIN>8 was considered acceptable). 10X Genomics Chromium microfluidic chips were loaded with around 9000 and cDNA libraries were generated following manufacturer’s protocol. Concentrations and fragment sizes were determined using Agilent Bioanalyzer and Invitrogen Qubit. cDNA libraries were sequenced using NextSeq 500 and High Output v2.5 (75 cycles) kits. Count matrices were subsequently generated following 10X Genomics Cell Ranger pipeline.

Following normalisation and quality control, we obtained an average of 5792 ± 1415 cells/condition.

### Seurat preprocessing

scRNAseq datasets were processed using Seurat (https://satijalab.org/seurat/) (Butler et al., 2018). Cells with more than 10% of mitochondrial gene fraction were discarded. 4000-5000 genes were detected on average across all 4 datasets. Dimensionality reduction and UMAPs were generated following Seurat workflow. The top 100 DEGs were determined using Seurat "FindAllMarkers" function with default parameters. When processed independently (scvelo), the datasets were first regressed on cell cycle genes, mitochondrial fraction, number of genes, number of UMI following Seurat dedicated vignette, and doublets were removed using DoubletFinder v3 (McGinnis et al., 2019). A "StressIndex" score was generated for each cell based on the list of stress genes previously reported (Machado et al., 2021) using the “AddModule” Seurat function. 94 out of 98 genes were detected in the combined datasets. UMAPs were generated after 1. StressIndex regression, and 2. after complete removal of the detected stress genes from the gene expression matrix before normalization. In both cases, the overall aspect of the UMAP did not change significantly (**Figure S5**). Although immeasurable confounding effects of cell stress following isolation cannot be ruled out, we reasoned that our datasets did not show a significant effect of stress with respect to the conclusions of our study.

### Matrisome analysis

After subsetting for the features of the Matrisome database (Naba et al., 2015) present in our single-cell dataset, the matrisome score was calculated by assessing the overall expression of its constituents using the "AddModuleScore" function from Seurat (Butler et al., 2018).

### RNA velocity and driver genes

Scvelo was used to calculate RNA velocities (Bergen et al., 2020). Unspliced and spliced transcript matrices were generated using velocyto (Manno et al., 2018) command line function. Seurat-generated filtering, annotations and cell-embeddings (UMAP, tSNE, PCA) were then added to the outputted objects. These datasets were then processed following scvelo online guide and documentation. Velocity was calculated based on the dynamical model (using *scv.tl.recover_dynamics(adata)*, and *scv.tl.velocity(adata, mode=’dynamical’)*) and differential kinetics calculations were added to the model (using *scv.tl.velocity(adata, diff_kinetics=True)*). Specific driver genes were identified by determining the top likelihood genes in the selected cluster. The lists of top 100 drivers for EOM and TA progenitors are given in Suppl Tables 10 and 11.

### Gene regulatory network inference and transcription factor modules

Gene regulatory networks were inferred using pySCENIC (Aibar et al., 2017; Sande et al., 2020). This algorithm regroups sets of correlated genes into regulons (i.e. a transcription factor and its targets) based on binding motifs and co-expression patterns. The top 35 regulons for each cluster was determined using scanpy "scanpy.tl.rank_genes_groups" function (method=t-test). Note that this function can yield less than 35 results depending on the cluster. UMAP and heatmap were generated using regulon AUC matrix (Area Under Curve) which refers to the activity level of each regulon in a given cell. Visualizations were performed using scanpy (Wolf et al., 2018). The outputted list of each regulon and their targets was subsequently used to create a transcription factor network. To do so, only genes that are regulons themselves were kept. This results in a visual representation where each node is an active transcription factor and each edge is an inferred regulation between 2 transcription factors. When placed in a force-directed environment, these nodes aggregate based on the number of shared edges. This operation greatly reduced the number of genes involved, while highlighting co-regulating transcriptional modules. Visualization of this network was performed in a force-directed graph using Gephi “Force-Atlas2” algorithm (https://gephi.org/). Of note, a force-directed graph is a type of visualization technique where nodes are positioned based on the principles of physics that assign forces among the set of edges and the set of nodes. Spring like attractive forces are used to attract pairs of edges towards each other (connected nodes) while repulsive forces, like those of electrically charged particles, are used to separate all pairs of nodes. In the equilibrium state for this system, the edges tend to have uniform length (because of the spring forces), and nodes that are not connected by an edge tend to be drawn further apart (because of the electrical repulsion).

### Gene ontology analysis

Gene ontology analyses were performed on top 100 markers (obtained from Seurat function FindAllMarkers) or on top 100 driver genes (obtained from scvelo), using Cluego (Bindea et al., 2009). “GO Molecular Pathway”, and “REACTOME pathways” were used independently to identify common and unique pathways involved in each dataset. In all analyses, an enrichment/depletion two-sided hypergeometric test was performed and p-values were corrected using the Bonferroni step down method. Only the pathways with a p-value lower than 0.05 were displayed.

### Literature scores

The scores for SMMCs (Giordani et al., 2019), FAPs (Oprescu et al., 2020), myotendinous junction B myonuclei (Kim et al., 2020), Twist2+ population(Liu et al., 2017), fetal MuSCs (Tierney et al., 2016), developing limb connective tissues (Lima et al., 2021) and the human skeletal muscle mesenchyme (Xi et al., 2020) were calculated by assessing the overall expression of of the markers of each population (Suppl Table 5) using the "AddModuleScore" function from Seurat (Butler et al., 2018).

### Microarray data and analysis

EOM, TA, Esophagus, Masseter and Diaphragm muscles from three adult TgPax7nGFP mice were digested and processed for FACs as described before. GFP+ cells were collected and RNA extracted using the RNeasy Micro Kit (Qiagen). After validation of the RNA quality with Bioanalyzer 2100 (using Agilent RNA6000 Pico Chip Kit), 450 pg of total RNA is reverse transcribed following the Ovation Pico WTA System V2 (Tecan). Briefly, the resulting double strand cDNA is used for amplification based on SPIA technology. After purification according to Tecan protocol, 3.6 ug of Sens Target DNA are fragmented and biotin labelled using Encore Biotin Module kit (Tecan). After control of fragmentation using Bioanalyzer 2100, cDNA is then hybridized to GeneChip® MouseGene2.0 (Affymetrix) at 45°C for 17 hours.

After overnight hybridization, chips are washed on the fluidic station FS450 following specific protocols (Affymetrix) and scanned using the GCS3000 7G. The scanned images are then analyzed with Expression Console software (Affymetrix) to obtain raw data (Cel files) and metrics for Quality Controls. Data preprocessing with the oligo R package: Cel files were imported in R with the read.celfiles() function. Background subtraction and quantile normalization were performed using the Robust Multichip Average algorithm (rma() function).The obtained expression matrix was then annotated using the getNetAffx() from the oligo library in combination with the pd.mogene.2.0.st annotation package. Low expressed genes were filtered out using the genefilter R packages functions; first the genes expression standard deviation was computed using the rowSds() function, then the shorth of the calculated standard deviation was computed and all genes with sd above the shorth were kept for the downstream analysis. Differential genes expression analysis was conducted with the Limma bioconductor package. Limma was used to fit a linear model with the pairwise combination of the different anatomical locations as a set of contrasts. This linear model was then used to compute the moderated t-statistics with the eBayes() function. Finally, the multiple test across genes and contrasts was performed using the decideTests() function with default parameters.

### Code availability

The code that was used to generate the TF network is available at this address: https://github.com/TajbakhshLab/TFnetwork

## Acknowledgements

We acknowledge funding support from the Institut Pasteur, Agence Nationale de la Recerche (Laboratoire d’Exellence Revive, Investissement d’Avenir; ANR-10-LABX-73 to ST and ANR-21-CE13-0005 MUSE to GC), Association Française contre les Myopathies (Grant #20510 to ST and #23201 to GC), and the Centre National de la Recherche Scientifique. M.B.D. was supported by a grant from Laboratoire d’Excellence Revive and La Ligue Contre le Cancer. We gratefully acknowledge the Genomic Patform (Inserm U1016-CNRS UMRS 8104 - Université de Paris Cité) at Institut Cochin, the UtechS Photonic BioImaging (Imagopole), C2RT, Institut Pasteur, supported by the French National Research Agency (France BioImaging; ANR-10–INSB–04; Investments for the Future) and the Center for Translational Science (CRT)-Cytometry and Biomarkers Unit of Technology and Service (CB UTechS) at Institut Pasteur for support in conducting this study.

## Author contributions

MBD, GC, and ST conceived the study. MDB, DDG and GC performed most of the experiments. PTL, SG, MM and BE contributed to in vitro experiments and image acquisition. AG, MBD and DDG performed sc-RNAseq. AG and MBD run the bioinformatic analysis with help from SM and VL. AG, MBD and GC wrote the manuscript with input from DDG and ST. GC and ST provided funding.

## Supplementary Information

### Supplementary Tables

**Table 1. Top 100 marker genes of EOM and TA of in vitro activated MuSCs**

**Table 2. Reactome Pathway Analysis for EOM and TA in vitro activated MuSCs**

**Table 3. Molecular Function Analysis for EOM and TA in vitro activated MuSCs**

**Table 4. List of top 100 marker genes of EOM and TA of quiescent MuSCs**

**Table 5. Genes used to generate signature scores for EOM and TA in vitro activated MuSCs**

**Table 6. List of top 100 regulons of Prog and Diff clusters of EOM and TA activated MuSCs**

**Table 7. Foxc1 targets identified by pyScenic**

**Table 8. List of top 100 regulons of the global EOM and TA activated MuSCs populations**

**Table 9. List of top 100 regulons of the global EOM and TA quiescent MuSCs populations**

**Table 10. List of top 100 drivers for EOM progenitors**

**Table 11. List of top 100 drivers for TA progenitors**

**Table 12. Antibodies and resources used in this study**

**Table 13. qRT-PCR primer list**

### Supplementary Figure Legends

**Suppl Figure 1. EOM and TA MuSCs exhibit functional differences following activation**

A. Experimental scheme. MuSCs were isolated by FACS based on GFP fluorescence from *Tg:Pax7-nGFP* mice. A fraction of these cells was individually plated on a 96-well plate (T1) and the remainder were activated for 2 days *in vitro* before re-sorting and clonal plating on a 96-well plate (T2). Cells were allowed to proliferate for 14 days and assessed for total cell number per clone.

B. Quantification of cell numbers per clone at both time points. Each dot represents a clone. Representative of 2 independent experiments. The mean of the clone size is indicated.

C. Western blots for AKT/Phospho-AKT, p38/Phospho-p38 and VINCULIN for normalization of protein loading. n =3 with cells coming from two mice pooled per lane.

D. Densitometric analysis of the Western Blots in D. Two-tailed unpaired Student’s t-test; *p-value<0.05.

**Suppl Figure 2. Single cell transcriptome signatures of activated EOM and TA MuSCs**

A. Quantification of Myod staining intensity. Violin plots show the single cell distribution Myod staining intensity of activated EOM and TA MuSCs from *Pax7^CreERT2^:R26^mTmG^*. Dots superimposed on violin plots correspond to the average value in the independent wells (n = 3 wells, each replicate encompasses measurements from >1000 cells). Grey bars represent mean ± SD. P values were calculated using a two-tailed unpaired Student’s *t* test (p-value < 2.2e-16).

B-C. Reactome pathway (A) and GO Molecular function (B) network analysis on top 100 DEGs of each cluster. Pie charts represent relative contribution of each cluster to this ontology term. Genes belonging to each term are highlighted in red.

**Suppl Figure 3. Molecular overlap of EOM and TA MuSCs across cell states**

A. Heatmap of adult quiescent MuSCs isolated from cranial mesoderm-derived (extraocular muscle (EOM), esophagus (ESO), masseter (MASS)) and somitic mesoderm derived (Tibialis anterior (TA), diaphragm (DIA)) muscles from *Tg:Pax7-nGFP* mice.

B. Heatmap from in vitro activated sc-RNAseq datasets of selected signature genes of EOM and TA MuSCs identified in Figure 3F,G.

C. Heatmap on quiescent sc-RNAseq datasets of selected signature genes of EOM and TA MuSCs identified in Figure 3F,G.

**Suppl Figure 4. EOM progenitors exhibit a distinctive mesenchymal cell signature**

A. Scores of EOM Prog, TA Prog, EOM Diff and TA Diff fractions for the signature of SMMCs (Giordani et al. Mol Cell 2019), FAPs and Fibroblasts (Oprescu et al. iScience 2020), myotendinous junction B myonuclei (Kim et al. Nat Com 2020), Twist2+ population (Liu et al. Nat Cell Biol 2017), fetal MuSCs (Tierney et al. Cell Reports 2016), developing limb connective tissues (Esteves-Lima et al. Nat Com 2021), Skeletal Muscle Mesenchyme (Xi et al. Cell Stem Cell 2020) and Global Myogenic markers.

B. Scheme of isolation of EOM and TA MuSCs from *Tg:Pax7-nGFP* mice, culture and staining for PDGFRβ-BV786 before flow cytometry analysis.

C. Representative flow cytometry profiles of EOM and TA activated MuSCs as per in E stained with PDGFRβ-BV786.

**Suppl Figure 5. Distinct gene regulatory networks underlie EOM and TA activation dynamics**

A. UMAP visualization of EOM and TA quiescent (top) and in vitro activated (bottom) clusters.

B. Expression plots of Foxc1, Creb3l1, Ebf1 and Hoxa10 in both datasets.

C. Scheme of isolation procedure, culture and Western Blot analysis (D-F) of EOM and TA MuSCs from *Tg:Pax7-nGFP* mice.

D. Western blots of EOM and TA MuSCs cultured for 4 days showing expression of FOXC1, EBF1, CREB3L1, PAX7, CYCLIND1, H3/TUBULIN for normalization of protein loading. Cells coming from n=3 mice pooled per lane.

E. Stress Index score across the merged activated sc-RNAseq dataset, overlaid onto UMAP representation.

F. Plot highlighting the correlation between Stress Index and the expression of myogenic markers *Pax7* (progenitor) and *Myod1* (committed/differentiating) on the activated sc-RNAseq dataset.

G. UMAP representations of different clusters and the expression of *Myod1* and *Pax7* before and after Stress Index regression, or removal of Stress Index genes from the count matrix. Upon regression the shape of the UMAPs is similar, although it comes out as a mirror image of the control.

**Suppl Figure 6. Increased expansion capacity of EOM MuSCs is retained after several passages in vitro**

A. FACS plots of *Tg:Pax7-nGFP;Myog^ntdTom^* EOM and TA MuSCs following 3 days in culture (Act), and one or two passages (P1, P2). Cells were isolated by FACS based on GFP fluorescence from *Tg:Pax7-nGFP;Myog^ntdTom^* and cultured *in vitro* for 5 days (Act) (n=4). A faction of cells was analysed by Flow cytometry while the rest of cells were plated for the next time point. This procedure was repeated for all time points (passages P1 and P2, n=3). Cells were split every 3 days.

B. Brightfield images of EOM and TA MuSCs (n=3) at indicated time points (D3, day3; D5, day5; passages P1 and P2). Cells were split every 3 days.

**Suppl Figure 7. EOM features emerge prenatally and are kept with passages in vitro**

A. Foetal (E18.5) and postnatal (P21) EOM and TA MuSCs from *Tg:Pax7-nGFP* mice were cultured for 48h then processed for RNA extraction. Graph shows RT-qPCR for *Myogenin* normalized to *Rpl13*. Two-tailed unpaired Student’s t-test; *p-value<0.05. n=4 mice.

B. FACS plots of *Tg:Pax7-nGFP;Myog^ntdTom^* EOM and TA MuSCs isolated from P7 mice.

C. Expression table for Foxc1 extracted from MuscleDB dataset (Terry et al. eLife 2018) showing higher expression of Foxc1 in EOM (depicted here as eye muscles) compared to 10 other muscle groups.

D. Scheme of siRNA treatment of EOM MuSCs isolated from adult *Tg:Pax7-nGFP* mice.

E. Quantification of experiment in E. At Day2, total cell numbers were counted and percentage of PAX7+ and FOXC1 + cells out of total number of cells in the well was assessed by immunostaining in scramble control (ctrl) and Foxc1 siRNA treated cells (n=3 mice).

F. qRT-PCR for *Myogenin* on control and Foxc1 siRNA treated EOM cells as per in E. Graph shows qRT-PCR results for *Myogenin* normalized to *Rpl13* on control and Foxc1 siRNA treated EOM cells as described in (n=3 mice).

**Suppl Figure 8. Cell cycle does not explain the velocity streams of in vitro activated EOM and TA MuSCs**

A. UMAP representation of EOM velocity streams overlaid with cluster and cell cycle information.

B. UMAP representation of TA velocity streams overlaid with cluster and cell cycle information.

