## Supplementary figures and images for "Extraocular muscle stem cells exhibit distinct cellular properties associated with non-muscle molecular signatures"

### Suppl Figure 1

SupplFigure 1 Benavente-Diaz et al.

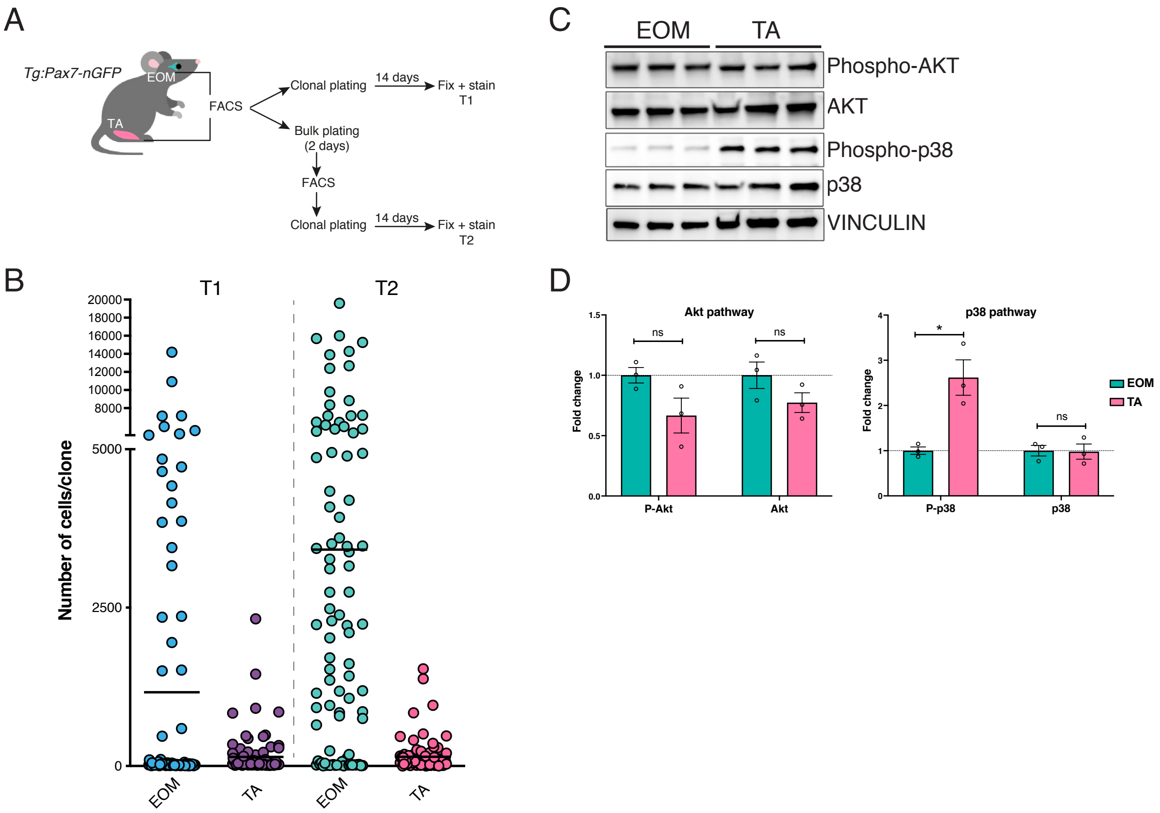

### Suppl Figure 2

Suppl Figure 2 Benavente-Diaz et al.

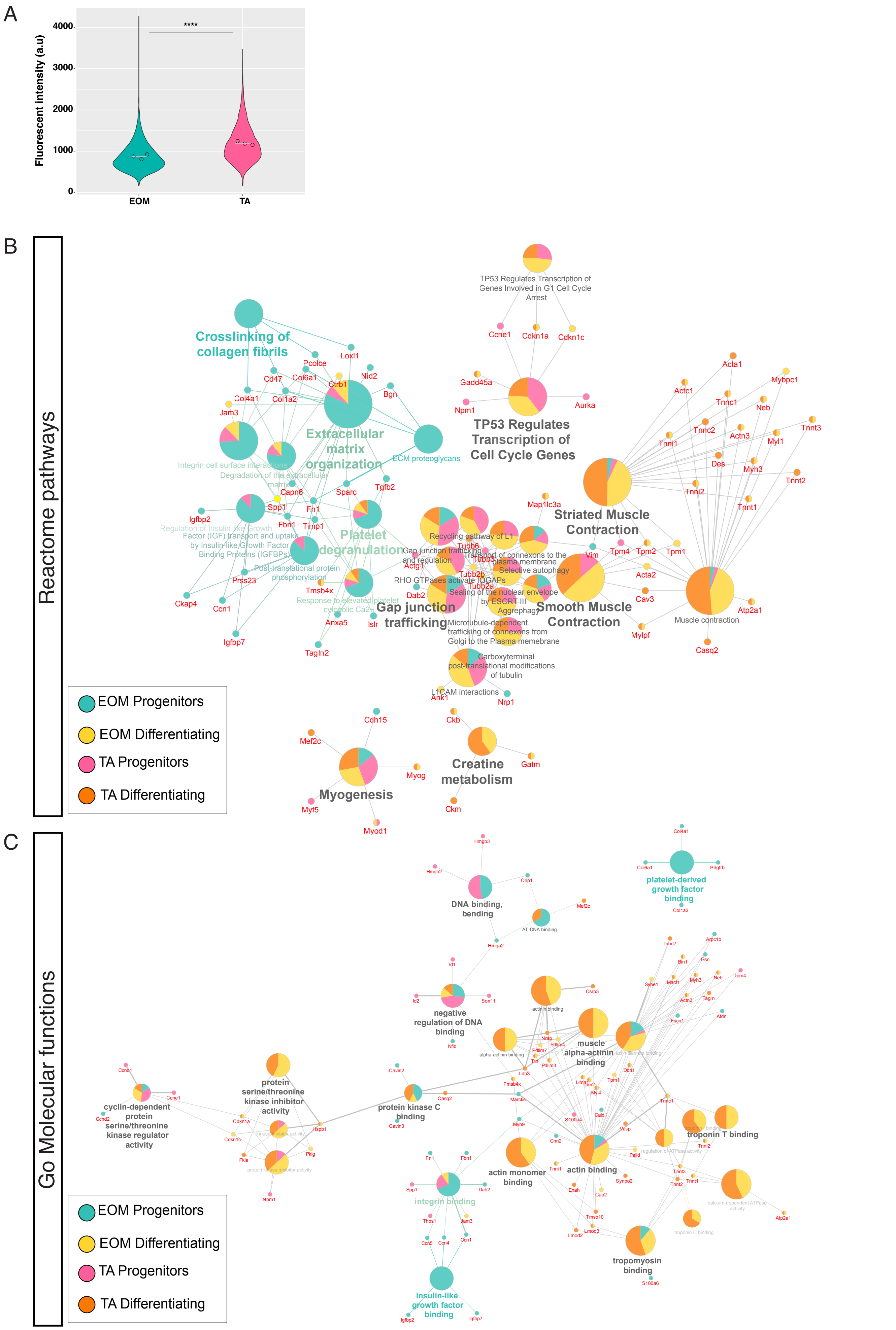

### Suppl Figure 3

SupplFigure 3 Benavente-Diaz et al.

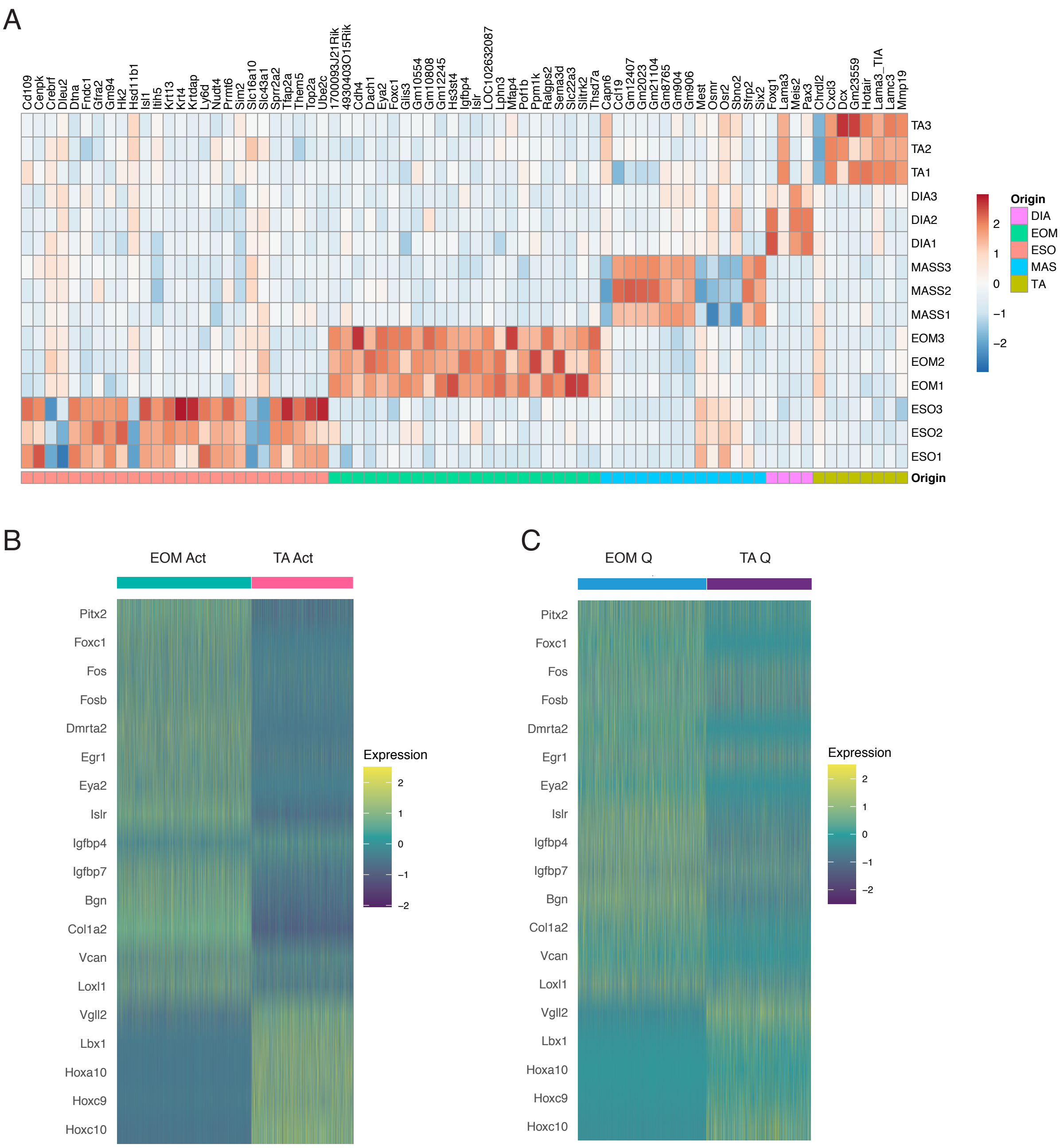

### Suppl Figure 4

A Litterature scores

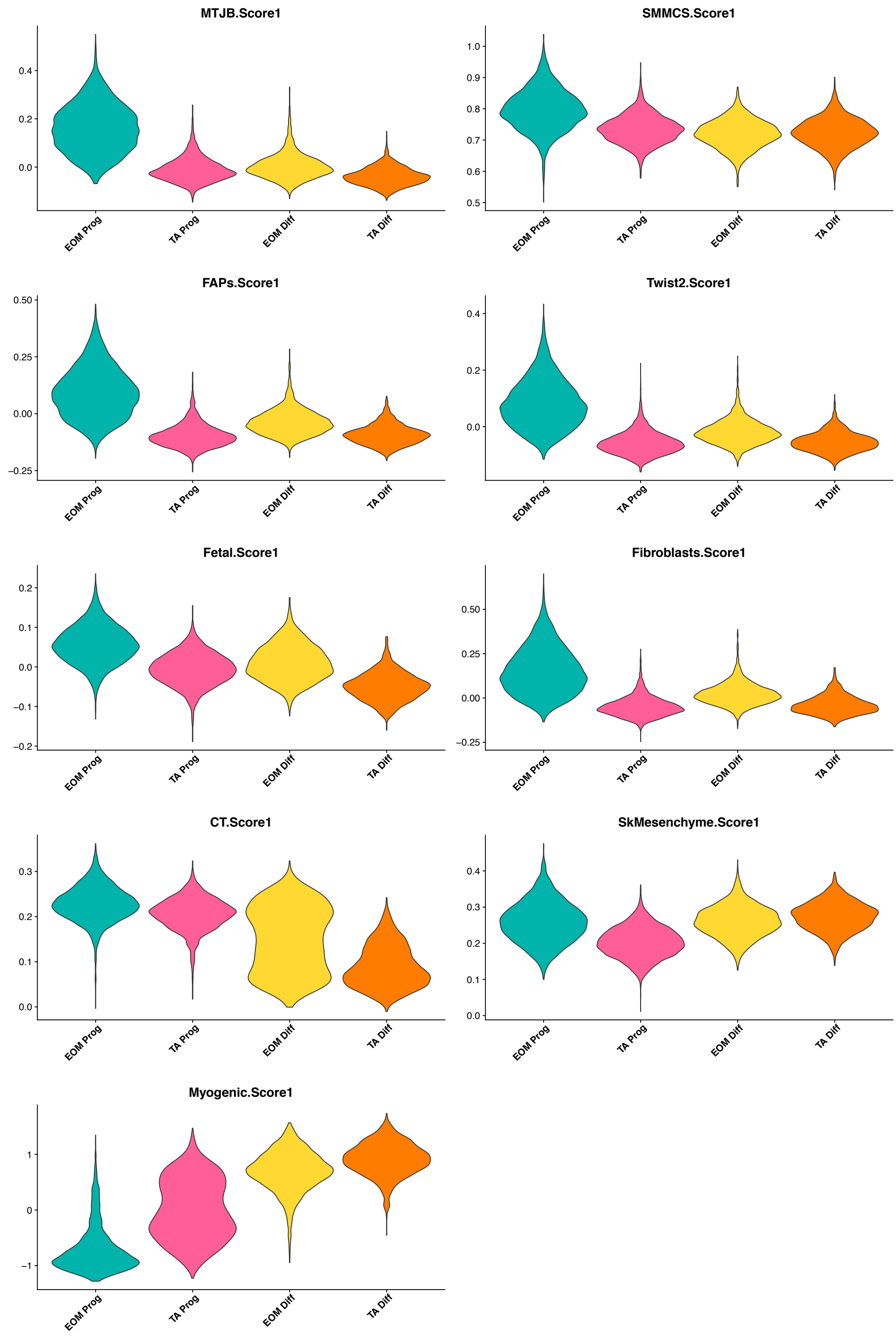


B

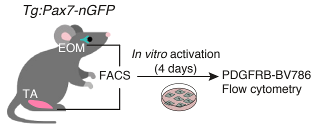


C

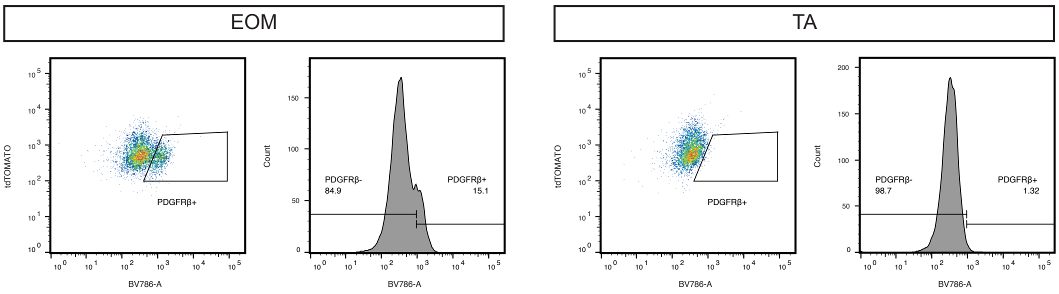

### Suppl Figure 5

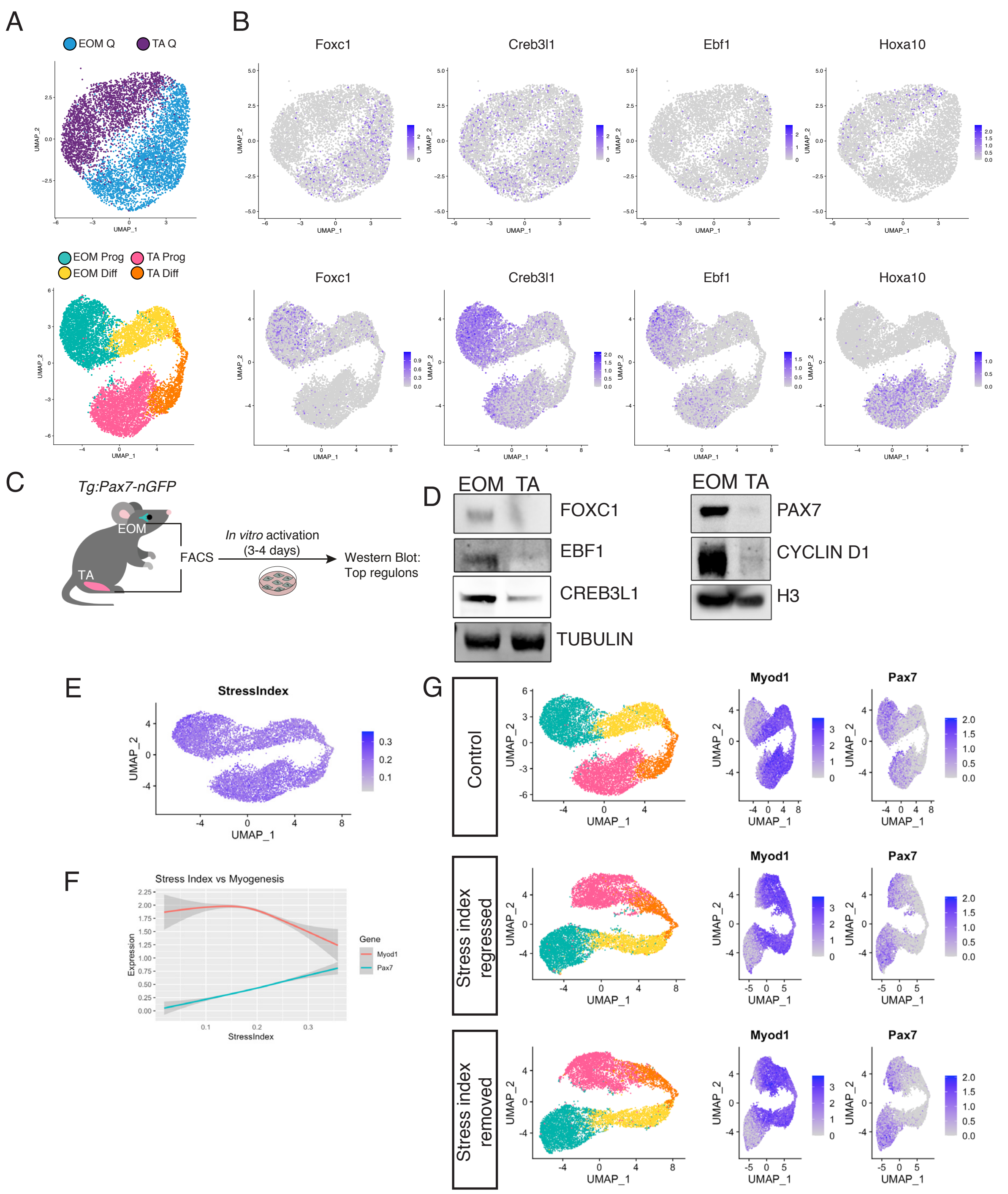

### Suppl Figure 5

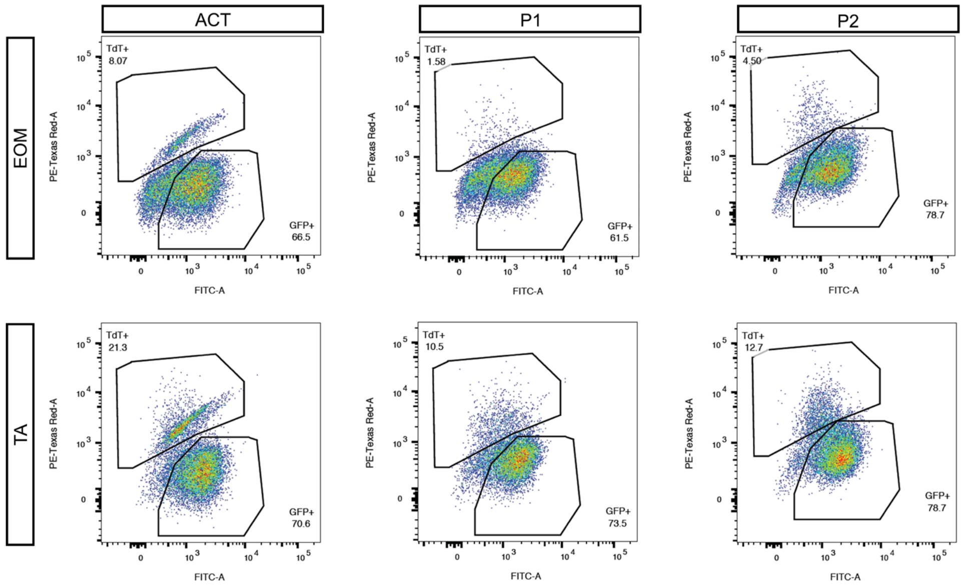

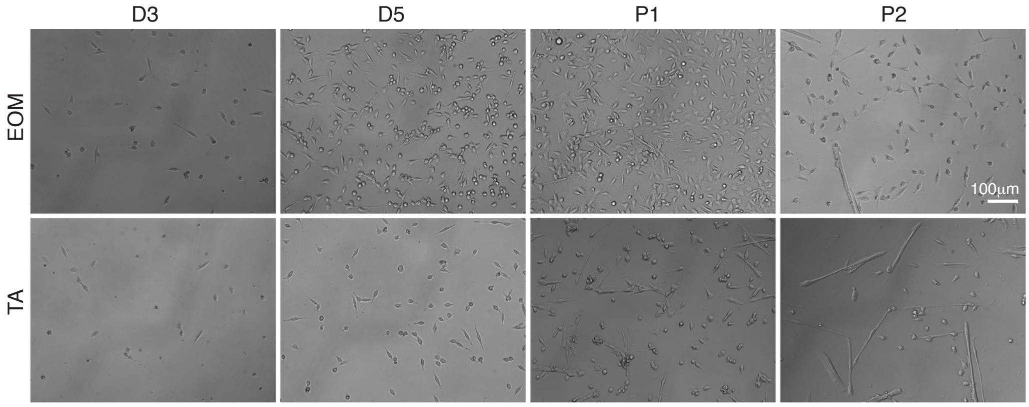

### Suppl Figure 7

SupplFigure 7 Benavente-Diaz et al.

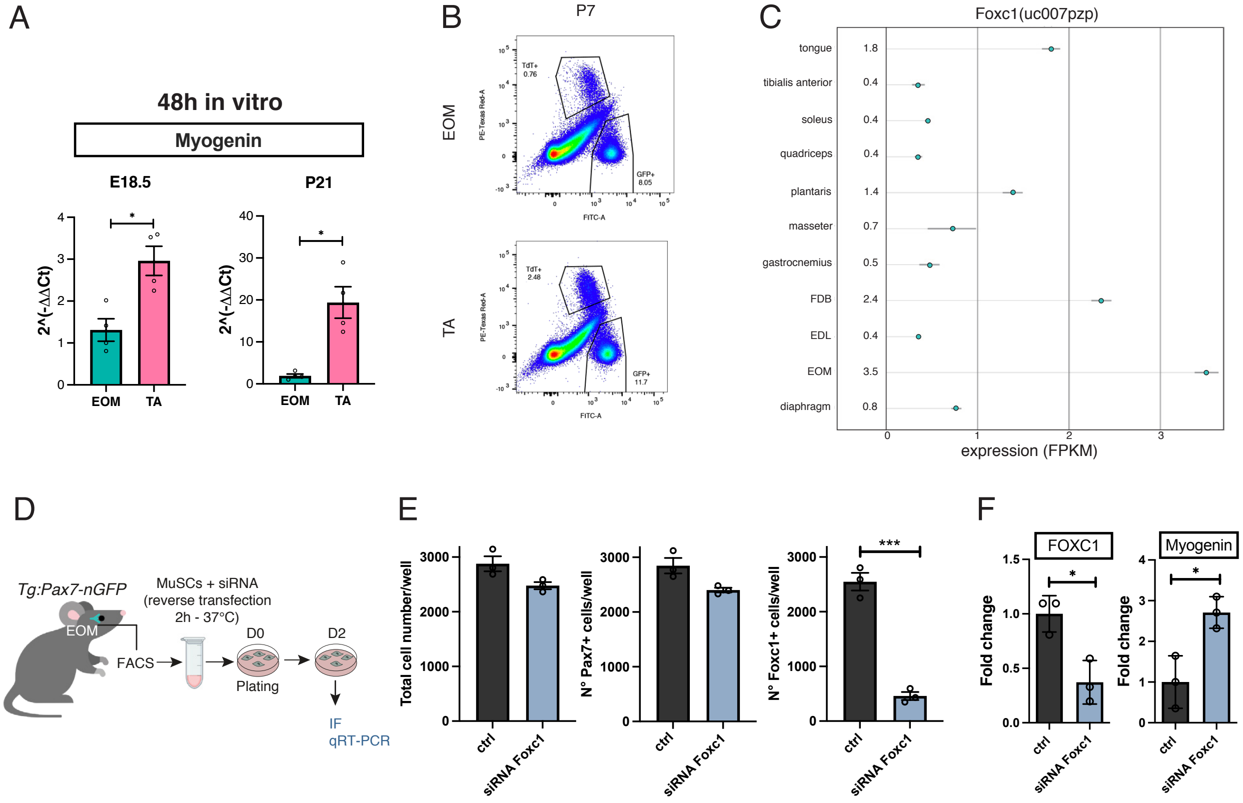

### Suppl Figure 8

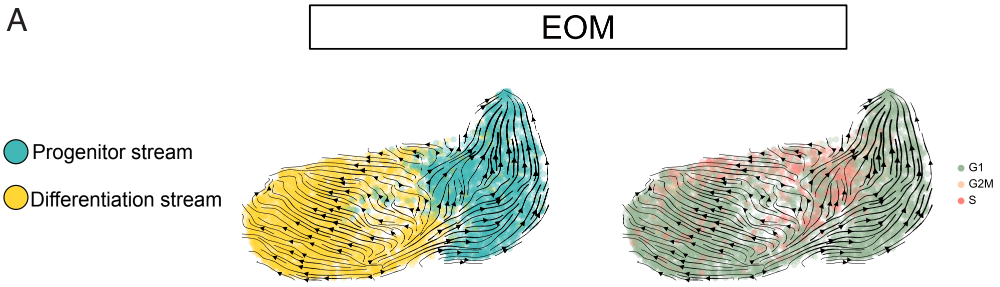

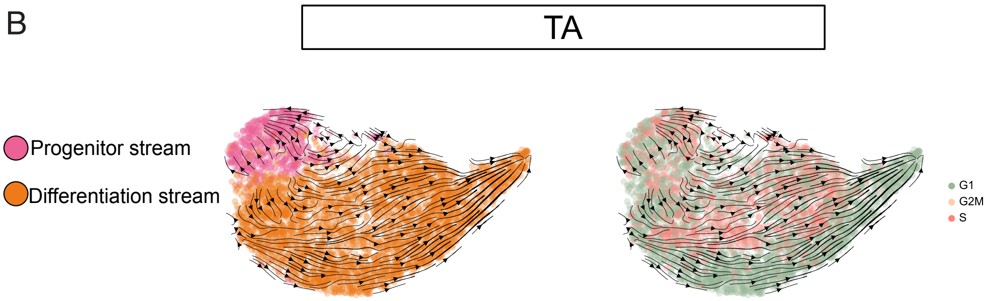
